# Rapid local adaptation in both sexual and asexual invasive populations of monkeyflowers (*Mimulus spp*.)

**DOI:** 10.1101/2020.12.19.423575

**Authors:** Violeta I. Simón-Porcar, Jose L. Silva, Mario Vallejo-Marín

## Abstract

**Background and Aims:** Traditionally, local adaptation has been seen as the outcome of a long evolutionary history, particularly in sexual lineages. In contrast, phenotypic plasticity has been thought to be most important during the initial stages of population establishment and in asexual species. We evaluated the roles of adaptive evolution and phenotypic plasticity in the invasive success of two closely related species of invasive monkeyflowers (*Mimulus*) in the United Kingdom (UK) that have contrasting reproductive strategies: *M. guttatus* combines sexual (seeds) and asexual (clonal growth) reproduction while *M. × robertsii* is entirely asexual.

**Methods:** We compared the clonality (number of stolons), floral and vegetative phenotype, and phenotypic plasticity of native (*M. guttatus*) and invasive (*M. guttatus* and *M*. × *robertsii*) populations grown in controlled environment chambers under the environmental conditions at each latitudinal extreme of the UK. The goal was to discern the roles of temperature and photoperiod on the expression of phenotypic traits. Next, we tested the existence of local adaptation in the two species within the invasive range with a reciprocal transplant experiment at two field sites in the latitudinal extremes of the UK, and analysed which phenotypic traits underlie potential local fitness advantage in each species.

**Key Results:** Populations of *M. guttatus* in the UK showed local adaptation through sexual function (fruit production), while *M*. × *robertsii* showed local adaptation via asexual function (stolon production). Phenotypic selection analyses revealed that different traits are associated with fitness in each species. Invasive and native populations of *M. guttatus* had similar phenotypic plasticity and clonality. *M*. × *robertsii* presents greater plasticity and clonality than native *M. guttatus*, but most populations have restricted clonality under the warm conditions of the south of UK.

**Conclusions:** Our study provides experimental evidence of local adaptation in a strictly asexual invasive species with high clonality and phenotypic plasticity. This indicates that even asexual taxa can rapidly (< 200 years) adapt to novel environmental conditions in which alternative strategies may not ensure the persistence of populations.

## Introduction

Populations of broadly distributed species adapt to local conditions through genetic differentiation (Williams, 1966; Kawecki and Ebert, 2004; Hereford, 2009) and phenotypic plasticity (Bradshaw, 1965; Donohue, 2013). These two mechanisms are universal, interacting, and non-mutually exclusive (Price *et al*., 2003; de Jong, 2005; West-Eberhard, 2005; Kelly, 2019). Yet, the traditional view was that local adaptation has a greater importance in sexual populations with a long evolutionary history (i.e. those with a greater number of recombination events behind; Weissmann, 1889; Crow and Kimura, 1965; Maynard Smith, 1968; Burt, 2000; Rushworth *et al*., 2020). In contrast, clonal propagation has been considered to reduce the opportunities for local adaptation (Schon *et al*., 1998; Rouzine *et al*., 2003; Schiffels *et al*., 2011) despite this mechanism can theoretically occur through selection on genes or genotypes (Vrijenhoek, 1979; Lushai *et al*., 2003). Given the expected reduction in genotypic diversity, phenotypic plasticity has been attributed a more important role in asexual lineages (Lynch, 1984; Van Kleunen and Fischer, 2001; Oplaat and Verhoeven, 2015; Fazlioglu and Bonser, 2016; Geng *et al*., 2016) and during the initial stages of population establishment (Davidson *et al*., 2011; Liao et al., 2016).

Introduced species often evolve to cope with novel biotic and abiotic conditions in non-native ranges (Bossdorf *et al*., 2005; Vandepitte *et al*., 2014; Oduor *et al*., 2016; Liu *et al*., 2020), and thus constitute an excellent model system to study adaptive evolution occurring over short periods of time (Thompson, 1998; Colautti and Lau, 2015). In addition, phenotypic plasticity seems to make a major contribution to the establishment and spread of introduced species in novel environments (Ghalambor *et al*., 2007; Riis *et al*., 2010; Ebeling *et al*., 2011; Pahl *et al*., 2013; Liao *et al*., 2016; Liu *et al*., 2016). Interestingly, clonal propagation is an advantageous trait for plant invasions, and numerous invasive plant species combine both sexual and asexual modes of reproduction or are mostly asexual (Pyšek, 1997; Silvertown, 2008; Roiloa, 2019). However, although an increasing number of studies have shown evolution at a contemporary scale in invasive plants with sexual (Lucek *et al*., 2004; Maron *et al*., 2004; Leger and Rice, 2007; Novy *et al*., 2013; Li *et al*., 2015; Bhattarai *et al*., 2017; Marchini *et al*., 2018) or mixed reproductive systems (Michel *et al*., 2004; Lambertini *et al*., 2010), field tests of local adaptation and phenotypic plasticity are rare for obligately asexual flowering plants (Mitchell and Whitney, 2018; Rushworth et al. 2020).

In this study, we investigate the evolutionary strategies of two invasive taxa that differ in their ability to reproduce sexually: *Mimulus guttatus* DC. and *M. × robertsii* Silverside (Phrymaceae). *Mimulus guttatus* (2n = 2x = 28) is an herbaceous, annual or perennial, plant native to Western North America (Grant, 1924; Wu *et al*., 2007; Lowry and Willis, 2010). *M. guttatus* was introduced in the United Kingdom (UK) for ornamental purposes 200 years ago (Roberts, 1964; Parker, 1975; Puzey and Vallejo-Marín, 2014) and perennial forms, which combine reproduction via seeds (sexual) and stolons (asexual), became naturalised in wetlands, riverbanks and wet ditches across the entire country (Preston *et al*., 2002), as in other areas in Europe and New Zealand (Howell and Sawyer, 2006; Truscott *et al*., 2006; Da Re *et al*., 2020). The second taxon, *Mimulus* × *robertsii*, is a triploid (2n = 3x = 44-46) originated in the UK, product of an unknown number of hybridisation events between introduced populations of the diploid *M. guttatus* and the closely related South American tetraploid *M. luteus* L. (2n = 4x = 60-62). *M. luteus* was introduced in the UK soon after *M. guttatus* but is currently rare (Vallejo-Marín and Lye, 2013). The hybrid *M* × *robertsii*, which is perennial and sexually sterile (Parker, 1975; Meeus *et al*., 2020) but capable of extensive clonal reproduction via stolons, has become well established across UK, though it is far less abundant than *M. guttatus* in the south range of the country (Preston *et al*., 2002; Stace, 2010; Vallejo-Marín and Lye, 2013; Da Re et al. 2020). *M. guttatus* and *M*. × *robertsii* are very similar in their morphology, phenology and habitat in the UK. Both species bear high genetic diversity and low genetic structure (Vallejo-Marín and Lye, 2013; Pantoja *et al*., 2017), suggesting that metapopulation dynamics with high gene flow are important in the spatial structuring of the introduced range.

We evaluate the roles of adaptive evolution and phenotypic plasticity in the invasive success of *Mimulus* at two nested levels: (i) between native and introduced populations, and (ii) among introduced populations. In a first experiment, we assess phenotypic differences and compared the clonality and plasticity of ancestral native (*M. guttatus*) and invasive (*M. guttatus* and *M*. × robertsii) populations under the environmental conditions at each latitudinal extreme of the UK, discerning the roles of temperature and photoperiod in a full-crossed design implemented in controlled environment chambers. Our hypothesis here is that asexual *M*. × *robertsii* should display levels of clonality and plasticity equal or higher than the sexual taxa (native *M. guttatus* and invasive *M. guttatus*). In a second experiment, we test the existence of local adaptation of the two species within the invasive range with a reciprocal transplant experiment at two field sites in the latitudinal ends of UK and analyse which phenotypic traits underlie the local fitness advantage in each species. We predict that if sexual reproduction increases the rate of adaptation, *M. guttatus* would be more likely to be locally adapted than *M. × robertsii*. To explore the possible mechanisms driving local adaptation, we also carry out phenotypic selection analyses to identify the phenotypic traits related to fitness in each species.

## Materials and Methods

### Experiment 1: controlled environment chambers

#### Plant material

For *M. guttatus*, we used seeds from five native populations from North America and from five introduced populations in the UK [Supplementary Information - Table S1]. We follow Lowry *et al*., (2019) and use the classic taxonomical definition of *M. guttatus* DC. (Grant, 1924), rather than the recent nomenclature proposed by Nesom (2014). All seeds were field-collected, except seeds from accessions in the Alaskan range, the putative ancestral range of UK populations (Puzey and Vallejo-Marín, 2014; Vallejo-Marin *et al*., 2020). Three Alaskan accessions were retrieved from herbarium specimens preserved at University of Alaska Museum Herbarium (accessions V153408, V127607, V142998). As each accession represents a single sampled individual and locality, these three accessions were pooled into a single Alaskan “population” (ALA). From each population, we selected three to five maternal seed families (seeds collected from the same maternal parent). In total, we had 43 families from 10 populations. For the sexually sterile hybrid *M*. × *robertsii*, we collected in the field vegetative fragments (clones) from five UK populations [Supplementary Information - Table S1]. In each population, we sampled 1-5 ramets (limited by population size) separated at least 1m to reduce the probability of sampling the same genet multiple times (15 ramets total from five populations). All maternal families in both species were randomly collected with regards of their clonality and phenotypic traits. Native and invasive populations of *M. guttatus* cover a wide latitudinal range of their distribution, while *M*. × *robertsii* populations proceeded from the centre and north of its narrower range.

#### Experimental treatments

We used the Controlled Environment Facility at the University of Stirling to create environmental conditions that resembled the UK *Mimulus* growing season (Fig. 1a). To model the conditions, we used two opposite localities that encompass the latitudinal range of *Mimulus* in the UK: Newport, in the Isle of Wight (50.70° N, 1.29° W), and Baltasound, in the Shetland Isles (60.76° N, 0.86° W). For each of these localities and for every two-week period between April and September (the UK *Mimulus* growing season), we calculated the photoperiod with the package *geosphere* (Hijmans, 2014) on *R* (R Core Team, 2019) and obtained maximum and minimum temperatures from the WorldClim database (Hijmans *et al*., 2005). Photoperiod and temperature temporal series were combined in a full-crossed design to create four experimental treatments that allowed us to disentangle the effects of temperature and photoperiod on plant performance: a short day, warm temperature treatment (SW; the natural conditions in Newport), a long day, cold temperature treatment (LC; the natural conditions in Baltasound), a short day, cool temperature treatment (SC) and a long day, warm temperature one (LW) [Supplementary Information - Fig. S1 and Table S2]. The different growth conditions in each temporal series were substituted every 10 days to allow completing the experiment in 120 days. Each experimental treatment was implemented in one Snijder Scientific (Tilburg, Netherlands) MC1750E controlled environmental chamber.

**Figure 1.**
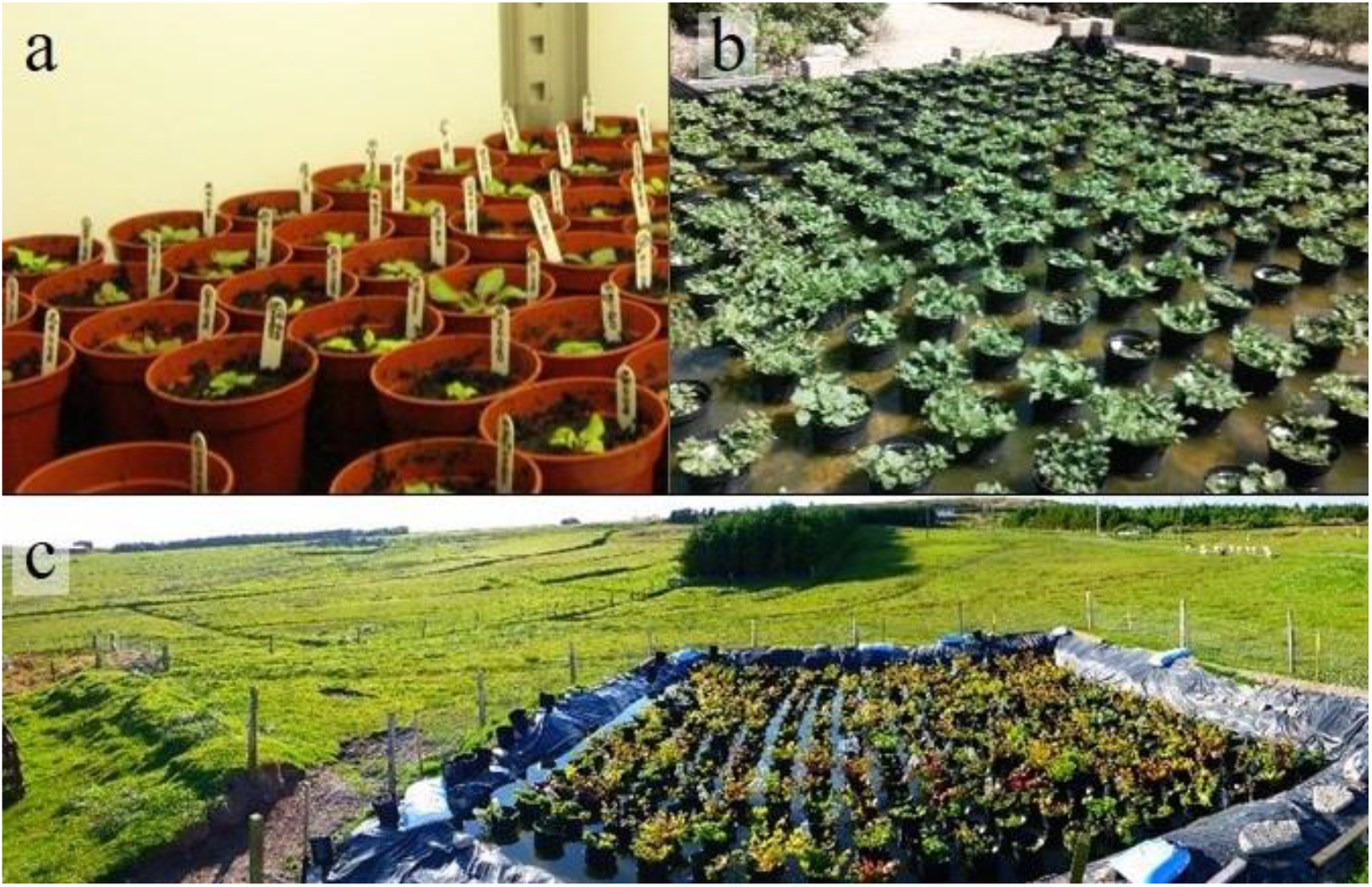
Experimental set ups to test for phenotypic plasticity and local adaptation of *Mimulus* in the UK. Seedlings growing in a controlled environmental chamber at the University of Stirling in 2014 (a). Common gardens set in the Isle of Wight (b) and Shetland (c) in summer 2015.

#### Plant growth

We planted seeds from each of 43 maternal families of *M. guttatus* into four 0.5 L pots (172 pots in total) filled with modular seed growing medium (Sinclair, Lincoln, Lincolnshire, UK), and placed them in the dark at 4°C for one week. For *M*. × *robertsii*, we planted individually eight cuttings from each of 15 maternal families in 0.5 L pots (120 pots in total) filled with All-Purpose growing medium (Sinclair, Lincoln, Lincolnshire, UK). All cuttings had a similar size, two small leaves and ∼2 cm. of roots. We moved one pot per maternal family of *M. guttatus* and two pots per maternal family of *M*. × *robertsii* to each chamber on 1^st^ May 2014. We noted the day of first germination for each *M. guttatus* pot and, four weeks after first germination, we selected and thinned the two biggest seedlings to one per pot filled with All-Purpose growing medium in order to get two replicates per maternal family in each chamber. The maximum difference in transplant time among pots was one week within each chamber and two weeks across the entire experiment. Pots were randomly repositioned within each chamber every other day throughout the experiment.

#### Measurements and statistical analyses

Most individuals survived until the end of the experiment, and all measurements were taken at this moment except otherwise specified. The experiment was stopped due to calendar constraints. To investigate the phenotypic differences in clonality among the three population types (classified by their origin and species, i.e., native *M. guttatus*, invasive *M. guttatus*, and *M*. × *robertsii*), we recorded the total number of stolons produced by individuals in the four environmental chambers. To compare their phenotypes, we recorded days to flower since germination or planting of the clonal fragment, corolla width of the first flower (measured with a digital calliper to the nearest 0.1mm in the second day after anthesis), whether plants flowered or not, the number of branches, floral stems and flowers, plant height (from the soil surface to the highest meristem, measured to the nearest cm), and length and diameter of the first internode. Finally, the entire individuals (above- and belowground) were harvested, washed out gently in water and dried at 60° C in individual paper bags for estimating final total dry biomass. In order to avoid over-parameterization in subsequent analyses, we averaged the two values from each family (cuttings in *M*. × *robertsii*, siblings in *M. guttatus*), for each trait under each of four treatments (except for germination time in *M. guttatus*, which had a single data point per family).

Preliminary analyses showed low correlation between most phenotypic traits [Supplementary Information - Fig. S2] and thus each variable was analysed separately. We used Generalized Linear Mixed Models (GLMMs) to analyse the variation in clonal reproduction and phenotypic traits as a function of the population type (native *M. guttatus*, introduced *M. guttatus* and *M*. × *robertsii*), treatment photoperiod (Short vs. Long), treatment temperature (Warm vs. Cold), and their two- and three-way interaction terms. Analogous GLMMs were also carried out for each population type separately. In all models, population was included as a random effect. We used a Poisson error distribution for number of stolons, branches, floral stems, and flowers, a binomial model for flowering, and a Gaussian model for germination time, flowering time, corolla width, plant height, internode length, internode diameter, and dry mass. The survival of plants was above 96% and thus this variable was not modelled. The significance of the fixed effects and their interactions were assessed by type-III Wald χ2 tests on the corresponding GLMMs. Where the interactions were not significant, we removed them and tested also the effect of the main effects alone with type-II Wald χ2 tests. To account for multiple tests, we applied a Bonferroni correction, dividing the significance alpha level by the number of variables analysed (corrected *P*-value = 0.004). Where population type or any interaction of fixed factors were significant, we performed *post hoc* contrasts based on estimated marginal means (EMMs) of the corresponding model. These procedures were repeated for all GLMMs in this study. All analyses were performed in R 3.4.0 (R Core Team, 2019) with packages lme4 (Bates *et al*., 2015), car (Fox *et al*., 2012) and emmeans (Lenth *et al*., 2018).

To investigate differences in phenotypic plasticity among population types, we estimated the Relative Distances Plasticity Index (RDPI; Valladares *et al*., 2006) for each trait measured in the chambers, for each family. RDPI were first estimated from trait distances between the two temperature (RDPI_*t*_) and the two photoperiod (RDPI_*p*_) treatments separately, pooling data from two chambers in each treatment. Trait distances were calculated as the absolute value of the difference of trait values of the same family (the average of the two individual replicates) at each of two treatments, divided by the maximum of the two trait values. RDPI were also estimated for each family across the four environmental chambers (RDPI_*tp*_) as the average of the six trait distances between each pair of chambers. We analysed the variation in phenotypic plasticity with GLMMs modelling RDPI_*t*_, RDPI_*p*_, and RDPI_*tp*_ estimates as a function of the population type. In addition, we run multivariate analyses of variance (MANOVAs) with RDPI_*t*_, RDPI_*p*_, or RDPI_*tp*_ estimates for all phenotypic traits as response variables and population type as independent variable. RDPI estimates for *germination day* were excluded for multivariate analyses because the lack of data for *M*. × robertsii. Finally, to test for differences in plasticity in response to temperature and photoperiod, we pooled RDPI_*t*_ and RDPI_*p*_ estimates and used a GLMM to model RDPI values as a function of the RDPI type, population type and their interaction. All RDPI GLMMs used a Gaussian distribution of errors.

### Experiment 2: Reciprocal transplants

#### Population survey and plant material

We used the distribution database of the Botanical Society of Britain and Ireland (BSBI; http://bsbidb.org.uk/) and personal records to design a survey of the northerb and the southern extremes of the distribution of *Mimulus guttatus* and *M*. × *robertsii* in the UK in summer 2014. We focused on BSBI records from the year 2000 with a precision of at least 100 m^2^. In total, we visited 60 localities between 50.1132° and 51.1489° N for the south of the country and 57.4963° and 60.8087° N for the north and found 39 populations. Because we were interested in identifying potential ecotypes adapted to the latitudinal extremes of the UK, we prioritized sampling fewer individuals in a higher number of populations instead of large numbers of individuals in fewer populations. This strategy has shown great statistical power (Blanquart *et al*., 2013) and was suited to our study system as many populations of *Mimulus* were small and likely contained few genets. To avoid sampling clones more than once, collected plants were at least 1m apart from each other. Cuttings were transported and planted at the greenhouse of the University of Stirling within one week after collection. In total, we sampled 155 cuttings from 36 populations for this study (Fig. 2) [Supplementary Information - Table S3]. *M. guttatus* and *M*. × *robertsii* are morphologically very similar and sometimes difficult to distinguish (Vallejo-Marín and Lye, 2013). To verify the species identity of each sampled individual we determined their relative genome size with flow cytometry (see methods in Simón-Porcar *et al*., 2017). To allow comparing M. × *robertsii* with both parental species, the only available population of *M. luteus* in the UK for which we had seeds was included in this experiment (Fig. 2) [Supplementary Information - Table S3]. For *M. luteus*, field-collected seeds from 25 different maternal individuals were planted and, once germinated, one seedling was transplanted and grown until adult.

**Figure 2.**
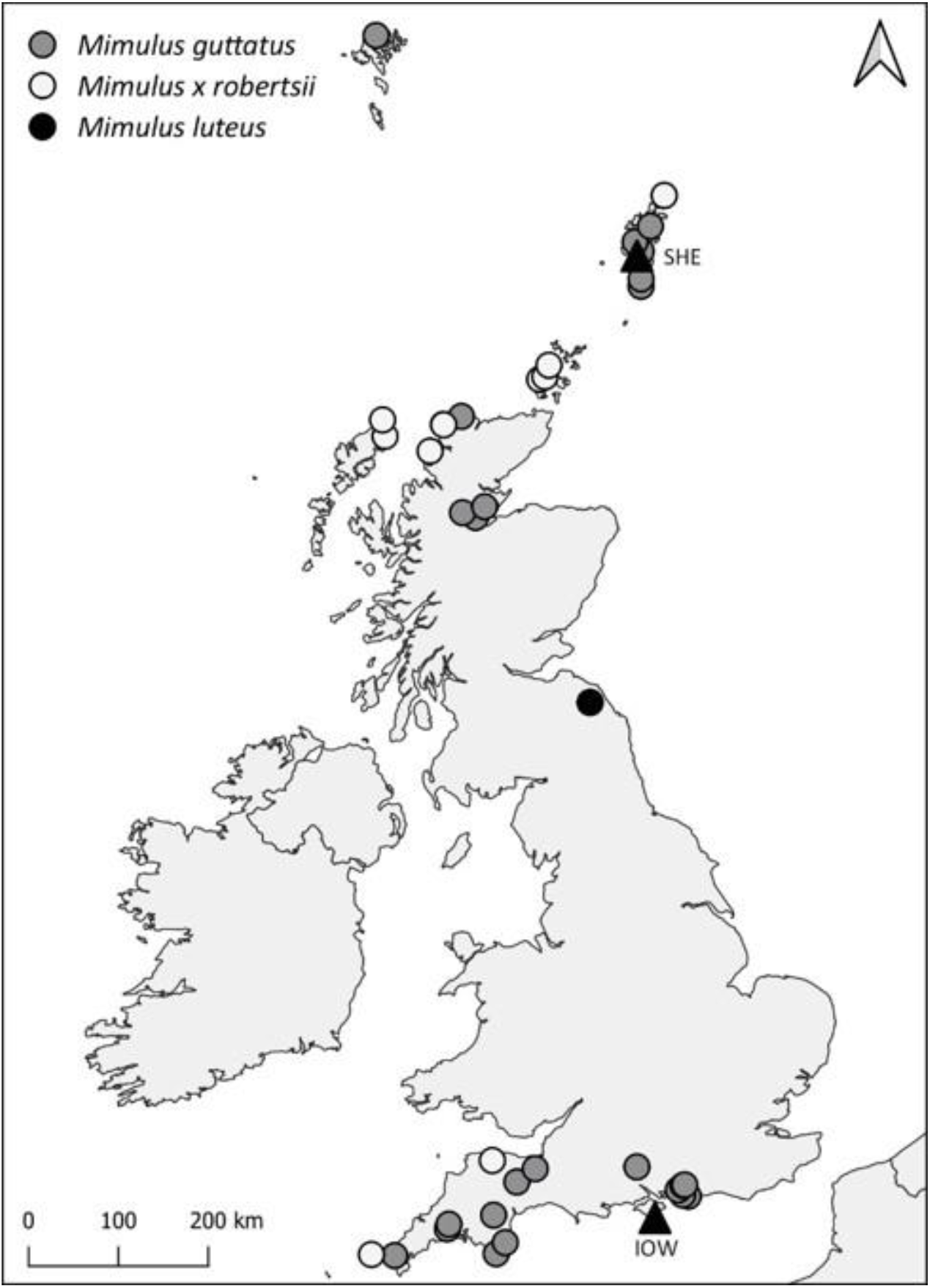
Map of populations (circles) and experimental sites (triangles) used in the reciprocal transplants.

We kept individual plants in 9-cm diameter pots filled with All-Purpose growing medium until next summer season. To minimize maternal resources effects, we transplanted each individual and randomized its position within the greenhouse at least four times between summer seasons. We cloned each individual four times to obtain replicates by April 2015, one month before setting up the experiment. All cuttings had similar architecture and size, and they were weighted prior to planting to evaluate possible individual size effects on subsequent measures of fitness. Clones were allowed to establish and develop roots and belowground biomass, similarly to how they naturally persist between growing seasons, but we restricted aerial biomass to the initial status by pruning elongating branches until the start of the experiment.

#### Experimental design

Two replicates per individual, for a total of 360 plants, were transplanted into each of two common gardens [Supplementary Information Table S3]. We established one common garden in the southernmost extreme of the UK at Ventnor Botanic Garden (Ventnor, Isle of Wight, England; 50.5890°, −1.2285°; IOW hereafter; Fig. 1b and 2) on May 14^th^ 2015, and a second common garden in the northernmost extreme of the UK at Da Gairdins i Sand (Sand, Shetland, Scotland; 60.2112°, −1.3761°; SHE hereafter; Fig. 1c and 2) on May 18^th^ 2015. The two common gardens were set up to be identical, consisting of a 100m^2^ square pond built up with a PVC pond liner (Aquatex, LBS Horticultural, UK), filled with 1cm of gravel to imitate natural conditions and provide an appropriate environment for root growth. Individual clones were planted in 10L pots filled with 7L of all-purpose commercial growing medium (LBS Horticultural, UK), which were placed in the pond in a regular grid with individuals from different species and origins completely randomized. Pots were 25 cm apart and pot walls precluded stolons to get out the pot, avoiding mingle or competition. The ponds were permanently flooded at a level of 10 cm so that plants were always moist as in natural habitats. The experiment was terminated at the end of the growing season, after senescence of the aerial parts of all individuals on August 24^th^ and 30^th^ in IOW and SHE, respectively.

#### Measurements and statistical analyses

To explore local adaptation, we assessed the fitness of individuals at each site recording their survival and reproductive success (number of stolons, and fruits in *M. guttatus*) at the end of the experiment. We also explored phenotypic differentiation and the traits contributing to local fitness differences within each species through phenotypic selection. For this aim we recorded plant height and stomata density (in the 6^th^ week of the experiment, when plants seemed to have achieved their maximum vigour); whether plants had flowered or not, plant cover, total dry biomass, and total number of branches, flowering stems, and flowers produced (at the end of the experiment); and days to flower, first flowering node, and corolla width of the first flower (at flowering of each individual). Stomata density, a trait involved in the hydric balance of plants and thus potential indicator of physiological variations (Raven 2002), was calculated under a 50X microscope from stomata imprints taken with transparent nail paint, adhesive tape and microscope slides from the beam of three new unshaded leaves per individual. Plant cover was measured over scaled overhead view photographs of each individual that were analysed with the software ImageJ (Abramoff *et al*., 2004). To estimate the dry biomass, the entire individuals (above- and belowground) were harvested, washed out and dried at 60°C in individual paper bags.

To ascertain the existence of local adaptation in *M. guttatus* and *M*. × *robertsii*, we analysed the variation in the sexual and asexual fitness measures (number of fruits and stolons) of each species with GLMMs, including Site, Origin, and their interaction as fixed factors in the models for each variable. The models used a Poisson distribution and included initial weight as covariate and population and individual nested within population as random factors. We consider a pattern of fitness advantage at home sites jointly with a significant effect of the interaction Site × Origin as evidence of local adaptation. The survival of plants was nearly 100% (see Results) and thus this variable was not modelled.

To explore the phenotypic differentiation between possible latitudinal ecotypes and compare the natural environmental effects on the growth of plants with the effects found in the environmental chambers experiment, we carried similar GLMMs for each species and phenotypic trait measured. A Poisson model was used to analyse the number of branches, floral stems, and flowers, and a Gaussian model was used for the remaining variables. We consider a significant effect of Origin as evidence of genetic differentiation, and a significant effect of Site as evidence of strong environmental effects (i.e. plasticity) on the development and growth of plants.

To investigate the phenotypic traits contributing to local fitness, we carried out phenotypic selection analyses by regressing the sexual and asexual fitness measures (number of fruits and stolons produced) on standardized phenotypic traits separately for each species and site. Preliminary analyses showed high correlation between various phenotypic traits for each species in this experiment [Supplementary Information - Fig. S3] and thus selection gradients were estimated to determine the magnitude and sign of directional and stabilizing selection on each trait, excluding indirect selection on correlated traits (Lande and Arnold, 1983). Separately for each species and site, we calculated the relative fitness (individual fitness divided by mean fitness) and standardized trait values (with a mean of 0 and a variance of 1). To improve the normality of the residuals in the regression models, the relative numbers of fruits and stolons were root squared. Quadratic regression coefficients were doubled to estimate the stabilizing/disruptive selection differentials (Stinchcombe *et al*., 2008).

To investigate the causes of the low occurrence of *M. luteus* in the UK and compare the patterns found in the hybrid *M*. × *robertsii* with both parental species, we assessed the sexual and asexual fitness and the phenotypic patterns of the single population included in our experiment. We recognise that the study of a single population does not allow robust inferences on the species patterns but given the great scarcity of *M. luteus* populations in UK, we still consider this approach worthy and valuable for species comparisons. The production of fruits and stolons, and each phenotypic trait measured, were analysed as a function of experimental site with GLMMs including initial weight as covariate and individual as random factor. Then, we compared the fitness of *M. luteus* with the other two *Mimulus* species and tested the phenotypic similarity of *M*. × *robertsii* and *M. luteus* with GLMMs including species, experimental site, and their interaction as fixed factors, initial weight as covariate, and population and individual nested within population as random factors. Because of *M. luteus* had a single population in the north, the southern populations of *M*. × *robertsii* and *M. guttatus*, and the variable “population origin” were excluded from these analyses. Finally, we carried out phenotypic selection analyses on *M. luteus* as explained above.

## Results

### Experiment 1: controlled environment chambers

#### Clonality

The population types differed in clonality (χ^2^ = 17.974; *P <* 0.001), with *M*. × robertsii producing the most stolons, significantly more than native *M. guttatus* (Fig. 3). Overall, clonal reproduction was not affected by photoperiod but it was affected by temperature (χ^2^ = 8.670; *P =* 0.003). The significant interaction of population type and temperature (χ^2^ = 32.035; P < 0.001) reflected that warm treatments increased clonality in both *M. guttatus* groups, but decreased clonality in *M*. × robertsii (Fig. 3) [Supplementary Information - Tables S4 and S5].

**Figure 3.**
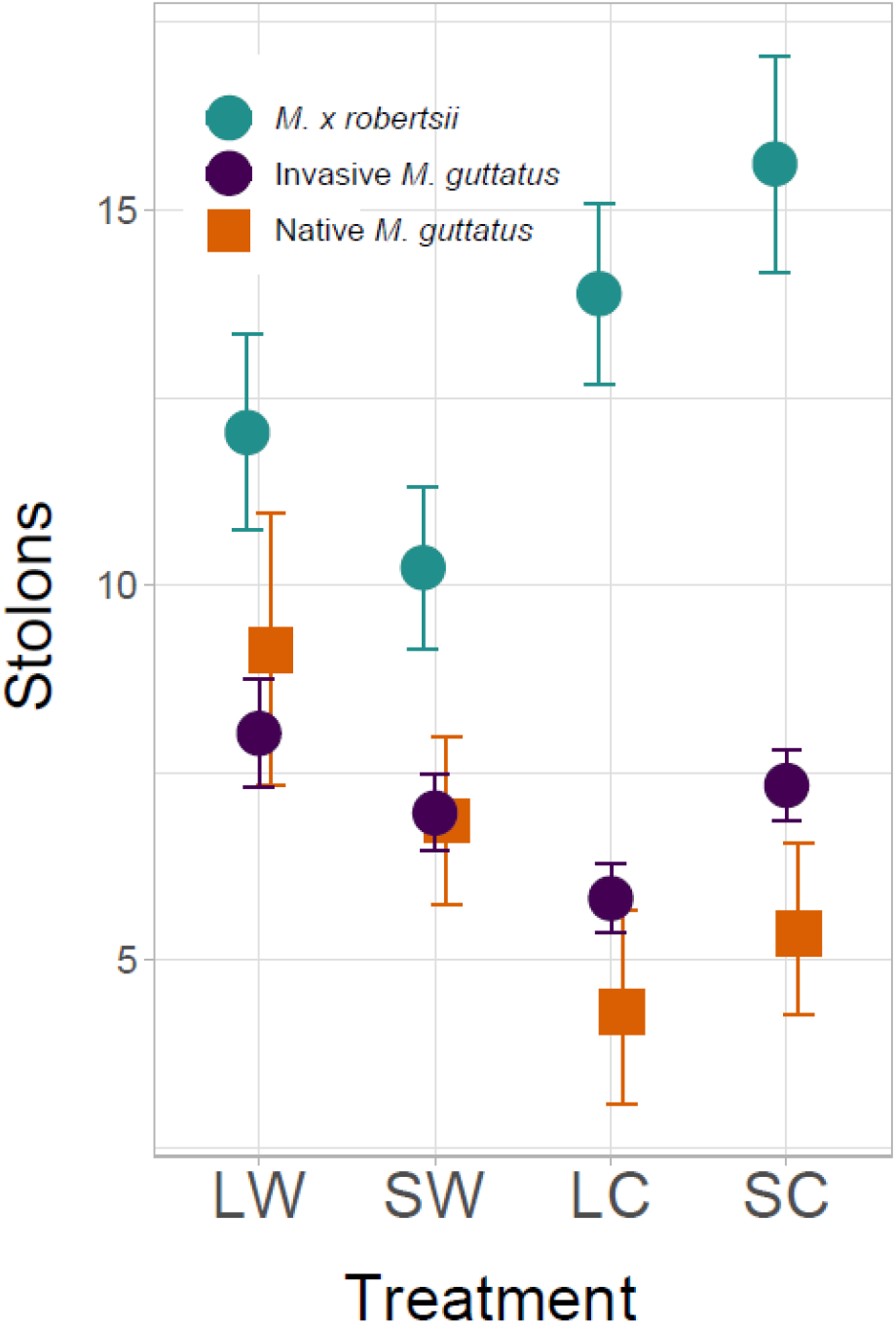
Clonality of *M*. × *robertsii* (ROB), introduced *M. guttatus* (UK) and native *Mimulus guttatus* (US) populations grown in four different controlled environmental chambers with contrasting photoperiods (L: long; S: short) and temperatures (C: cold; W: warm) in a crossed design. Mean values and standard errors of the variables measured are indicated by dots and error bars, respectively.

#### Phenotypes

Overall, *M*. × robertsii plants were shorter, thinner, and produced fewer branches, floral stems and flowers than both *M. guttatus* groups. Invasive *M. guttatus* produced fewer floral stems and flowers than native *M. guttatus* (χ^2^ > 13.928; *P <* 0.001) [Supplementary Information - Fig. S4 and Tables S4 and S5]. Warm treatments strongly accelerated germination and flowering in all population types, increased flower production, most significantly in native *M. guttatus*, and decreased corolla width, most significantly in invasive *M. guttatus*. Warm treatments also increased plant height in all groups, more sharply in *M. guttatus* than in *M*. × robertsii, increased internode length and decreased internode diameter and dry mass, most significantly in invasive *M. guttatus*, and increased the number of branches in both *M. guttatus* groups (χ^2^ > 9.094; *P <* 0.002) [Supplementary Information - Fig. S4 and Tables S4 and S5]. Short days delayed flowering in both *M. guttatus* groups, and strongly decreased the probability of flowering in invasive *M. guttatus* and *M*. × *robertsii*, the production of stems in *M. guttatus*, and flower production in all groups, more markedly in *M. guttatus* than in *M*. × *robertsii*. Short days also reduced plant height, most significantly in invasive *M. guttatus* and *M*. × *robertsii*, decreased internode length in both *M. guttatus* groups, and decreased internode diameter and dry mass, most significantly in invasive *M. guttatus* (χ^2^ > 8.911; *P <* 0.002) [Supplementary Information - Fig. S4 and Tables S4 and S5]. The interaction of temperature and photoperiod had an effect on the production of flowers in *M. guttatus*, with SC and LW treatments having the lowest and greatest flower production, respectively (χ^2^ > 14.412; *P* < 0.001). The three-way interaction of factors was always non-significant [Supplementary Information - Fig. S4 and Tables S4 and S5].

#### Phenotypic plasticity

The overall values for RDPI_*t*_, RDPI_*p*_ and RDPI_*tp*_ were 0.307 ± 0.031, 0.27 ± 0.034 and 0.32 ± 0.028 (mean ± sd), respectively. RDPI_*t*_ estimates did not differ among groups for any trait after Bonferroni correction (χ^2^ < 8.678; *P >* 0.013). The RDPI_*p*_ estimates for flowering day, number of flowers, floral stems, branches and plant height varied significantly among population types (χ^2^ > 11.991; *P <* 0.002). In most cases the *post hoc* tests indicated significantly greater plasticity in *M*. × robertsii than in the other groups [Supplementary Information - Table S6]. RDPI_*tp*_ estimates for dry mass were also significantly greater in *M*. × robertsii than in the other groups (χ^2^ = 13.655; *P =* 0.001) [Supplementary Information - Table S6]. MANOVAs found significant differences among population types for RDPI_*tp*_, RDPI_*t*_, and RDPI_*p*_ estimates (Pillai’s trace = 0.642-0.903; *F* > 2.261; *P* < 0.01; Table 1). *M*. × robertsii showed the greatest RDPI values, although the *post hoc* analyses showed only significantly differences in RDPI_*p*_ between *M*. × robertsii and native *M. guttatus* (Table 1). In the GLMM pooling RDPI_*t*_ and RDPI_*p*_ estimates, all fixed factors (RDPI type, population type and their interaction) were significant (χ^2^ > 6.716; *P <* 0.02). *M. guttatus* had higher RDPI_*t*_ than RDPI_*p*_ estimates and the opposite was found in *M*. × robertsii (differences were significant only within native *M. guttatus*). RDPI_*p*_ estimates of *M*. × robertsii were significantly higher than RDPI_*t*_ estimates of native *M. guttatus*. All RDPI estimates for the production of stolons were similar for all population types (χ^2^ < 5.034; *P >* 0.081) [Supplementary Information - Table S6].

**Table 1.** Comparison of phenotypic plasticity among population types. Results of the MANOVAs and *post hoc* tests analysing RDPI phenotypic plasticity indexes for all traits measured in the controlled environmental chambers as a function of the population type. Mean ± s.d. RDPI values across traits for each population type are provided jointly with the results of the post hoc tests. Different letters indicate significant differences. * P<0.05; ** P<0.01; *** P<0.001.

| | MANOVA | | | | | | | Mean ± sd; *post hoc* | | |
| --- | --- | --- | --- | --- | --- | --- | --- | --- | --- | --- |
|  | Df | df<br>residuals | Pillai's<br>trace | approx<br>F | num<br>df | den<br>df | *P* | *M. guttatus*<br>native | *M. guttatus*<br>invasive | *M. × robertsii* |
| RDPI<sub>t</sub> | 2 | 50 | 0.642 | 2.261 | 18 | 86 | 0.006 ** | 0.312 ± 0.052 a | 0.291 ± 0.042 a | 0.318 ± 0.071 a |
| RDPI<sub>p</sub> | 2 | 47 | 0.830 | 3.154 | 18 | 80 | < 0.001 *** | 0.218 ± 0.041 a | 0.247 ± 0.056 ab | 0.354 ± 0.078 b |
| RDPI<sub>tp</sub> | 2 | 32 | 0.903 | 2.287 | 18 | 50 | 0.011 * | 0.299 ± 0.038 a | 0.317 ± 0.046 a | 0.346 ± 0.065 a |

### Experiment 2: reciprocal transplants

#### Local adaptation

The survival and flowering of plants was respectively above 98% and 96% across the experiment. The fruit set of *M. guttatus* populations was significantly dependent on the experimental site and the interaction of experimental site and population origin (χ^2^ > 50.669; *P <* 0.001; Table 2). This species produced less fruits in SHE than in IOW, with a significant higher decrease for southern populations (Fig. 4). The production of stolons was not significantly dependent on any modelled factor in *M. guttatus* (Table 2), but it was also dependent on the experimental site and the interaction of experimental site and population origin in *M*. × robertsii (χ^2^ > 14.81; *P <* 0.001; Table 2). Overall, the production of stolons in *M*. × robertsii was higher in SHE than in IOW, and this was based on a high increase in northern populations. On the contrary, southern populations showed a slightly lower production of stolons in SHE than in IOW (Fig. 4).

**Table 2.**
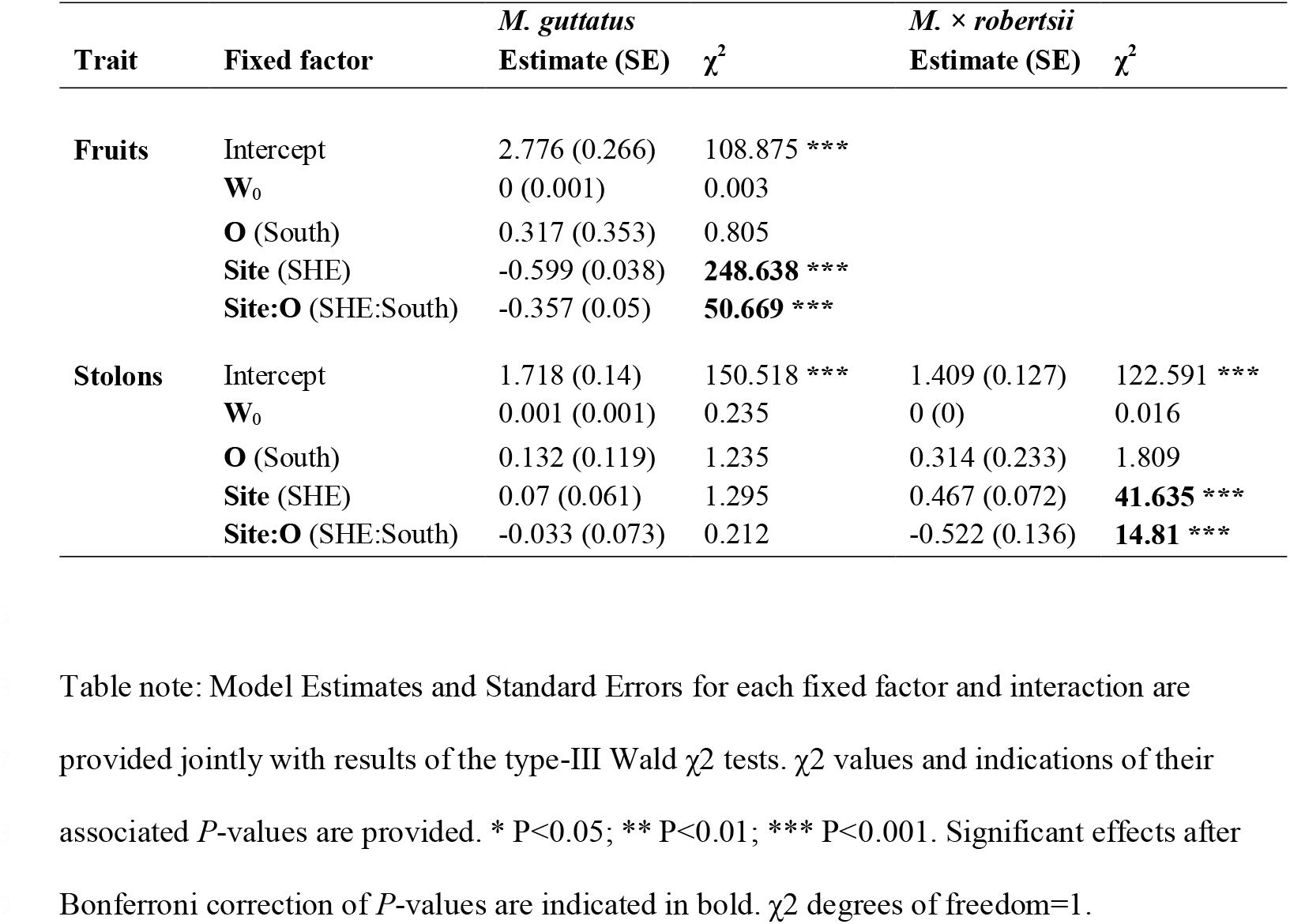
Results of the GLMMs modelling the effects of experimental site (S), population origin (O) and their interaction in the sexual and asexual fitness traits recorded in a reciprocal transplant experiment with introduced *Mimulus guttatus* and *M*. × *robertsii* populations.

| Trait | Fixed factor | *M. guttatus* |  | *M. × robertsii* |  |
| --- | --- | --- | --- | --- | --- |
| | | Estimate (SE) | $\chi^2$ | Estimate (SE) | $\chi^2$ |
| **Fruits** | Intercept | 2.776 (0.266) | 108.875 *** |  |  |
|  | **W<sub>0</sub>** | 0 (0.001) | 0.003 |  |  |
|  | **O** (South) | 0.317 (0.353) | 0.805 |  |  |
|  | **Site** (SHE) | -0.599 (0.038) | **248.638 \*\*\*** |  |  |
|  | **Site:O** (SHE:South) | -0.357 (0.05) | **50.669 \*\*\*** |  |  |
| **Stolons** | Intercept | 1.718 (0.14) | 150.518 *** | 1.409 (0.127) | 122.591 *** |
|  | **W<sub>0</sub>** | 0.001 (0.001) | 0.235 | 0 (0) | 0.016 |
|  | **O** (South) | 0.132 (0.119) | 1.235 | 0.314 (0.233) | 1.809 |
|  | **Site** (SHE) | 0.07 (0.061) | 1.295 | 0.467 (0.072) | **41.635 \*\*\*** |
|  | **Site:O** (SHE:South) | -0.033 (0.073) | 0.212 | -0.522 (0.136) | **14.81 \*\*\*** |

Table note: Model Estimates and Standard Errors for each fixed factor and interaction are provided jointly with results of the type-III Wald $\chi^2$ tests. $\chi^2$ values and indications of their associated *P*-values are provided. \* *P*<0.05; \*\* *P*<0.01; \*\*\* *P*<0.001. Significant effects after Bonferroni correction of *P*-values are indicated in bold. $\chi^2$ degrees of freedom=1.

**Figure 4.**
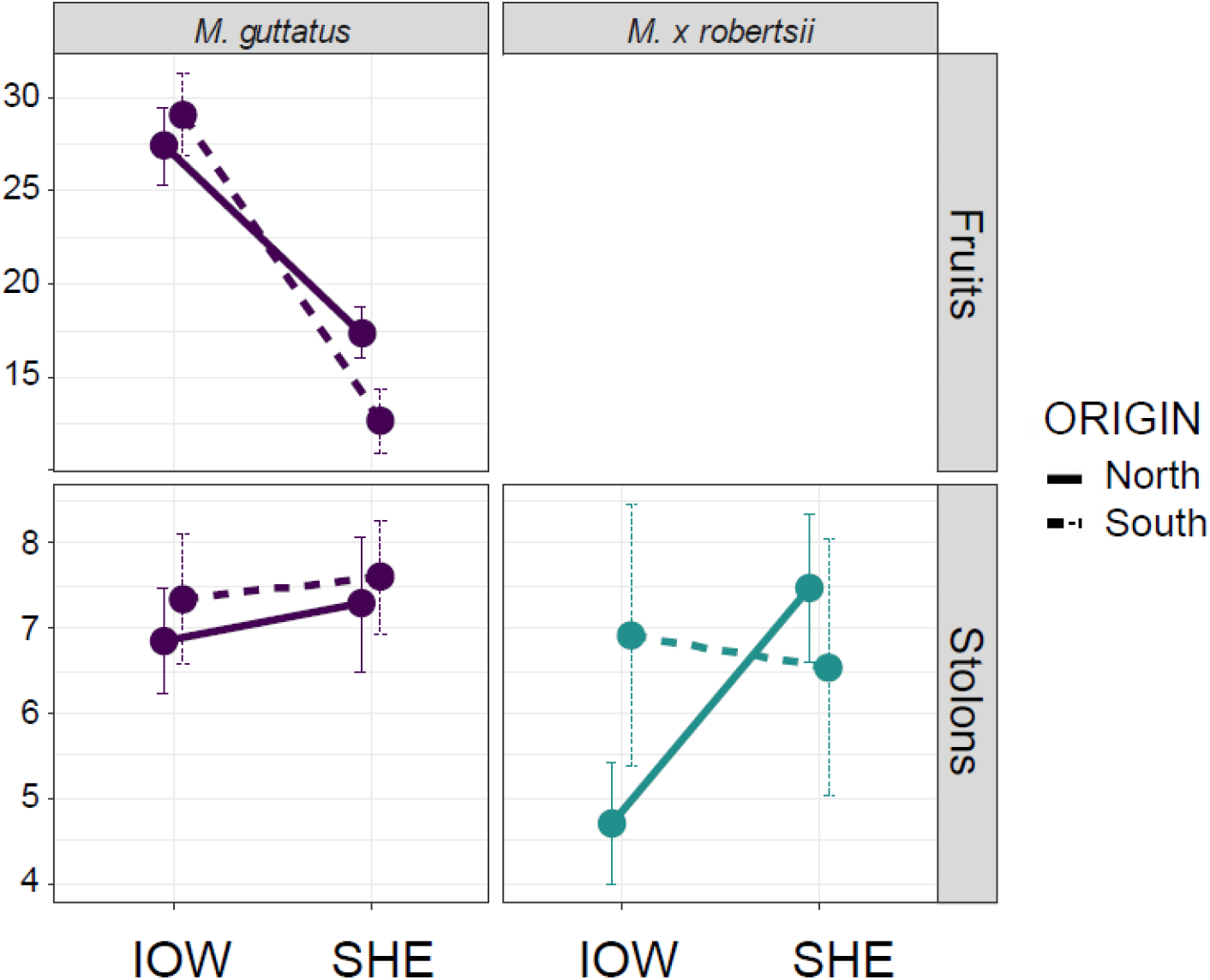
Fitness of *M. guttatus* and *M*. × robertsii individuals included in the reciprocal transplant experiment between different latitudes in the UK. Mean values and standard errors of the number of fruits and stolons are indicated by dots and error bars, respectively.

#### Phenotypic differentiation

Across most traits and for both species, plants were similar regardless of their latitudinal origin. In *M. guttatus*, northern individuals produced flowers with bigger corollas than southern individuals (χ^2^ = 6.566; *P* = 0.01) [Supplementary Information - Table S7 and Fig. S5]. Experimental site had a strong effect on the development of plants. *M. guttatus* individuals flowered later, produced less flowers, had lower stomata density and grew less according to plant cover and final dry mass in SHE than in IOW (χ^2^ > 24.524; *P* < 0.001). *M*. × *robertsii* flowered later, produced fewer floral stems and flowers, had lower dry mass, and produced more stolons, in SHE than in IOW (χ^2^ > 8.4; *P* < 0.01) [Supplementary Information - Table S7 and Fig. S5]. The interaction site x origin was significant for the number of branches, stems and flowers in *M. guttatus*, with negative estimates for south plants in SHE. In *M*. × *robertsii*, site x origin was significant for the number of flowers, with positive estimates for south plants in SHE (χ^2^ > 9.933; *P* < 0.01) [Supplementary Information - Table S7 and Fig. S5].

#### Phenotypic selection

The selection gradients differed between species, fitness traits and sites, suggesting diverse mechanisms of local adaptation in each species. In *M. guttatus*, fruit set (sexual fitness) in IOW showed significant positive linear selection and stabilizing selection on flowering day and dry mass, and significant negative linear selection and disruptive selection in height (*t* > 2.453; *P* < 0.015). In SHE there was significant positive linear selection in corolla width (*t* = 2.305; *P* = 0.023; Fig. 5) [Supplementary Information - Table S8]. For *M. guttatus* stolons (asexual fitness) we found only significant negative linear selection and disruptive selection in dry mass and number of branches in SHE (*t* > 2.12; *P* < 0.036; Fig. 5). For *M*. × *robertsii* stolons we found positive linear selection in dry mass in IOW, and significant positive linear selection and stabilizing selection in corolla width in SHE (*t* > 2.073; *P* < 0.042; Fig. 5) [Supplementary Information - Table S8].

**Figure 5.**
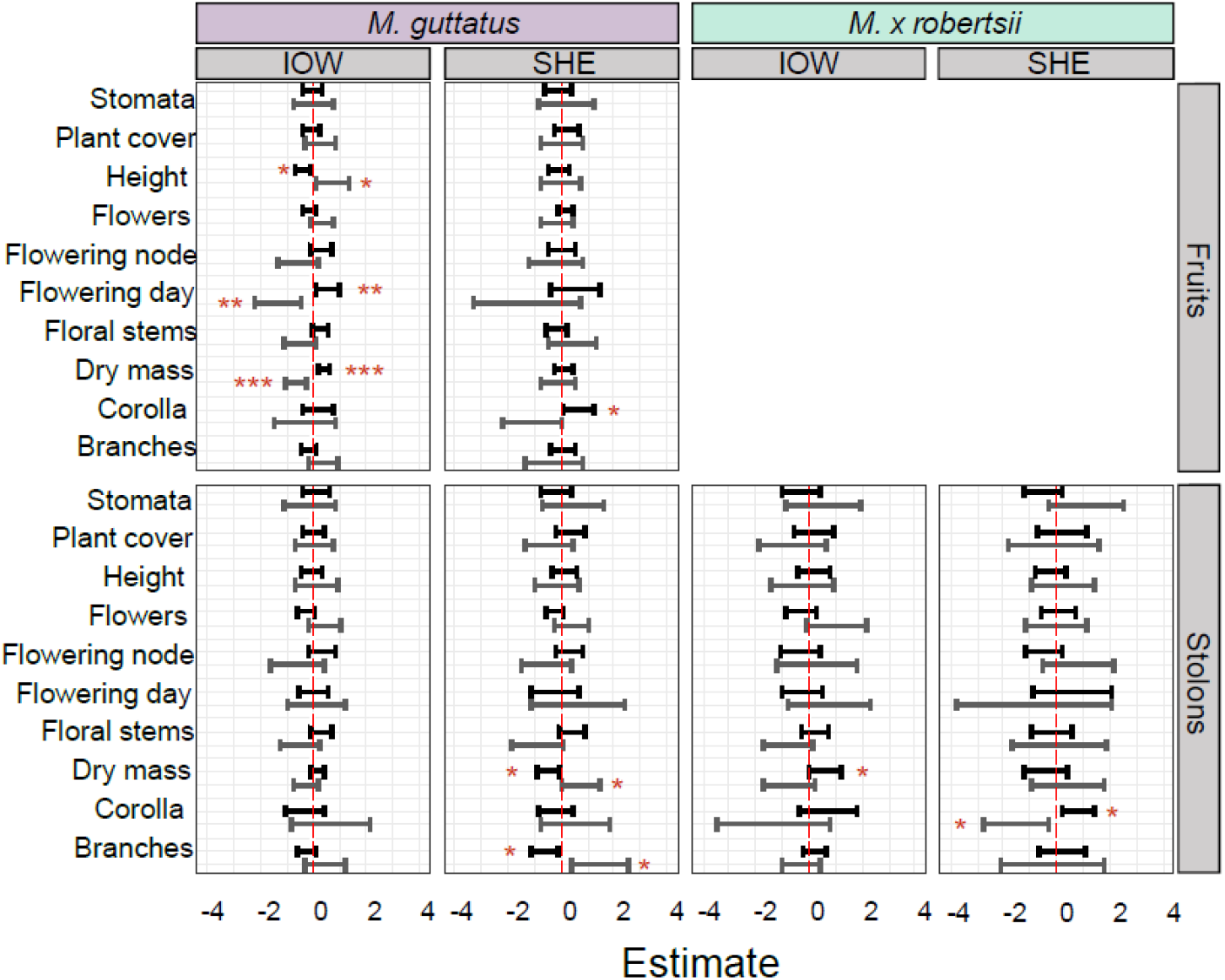
Estimates and 95% confidence intervals for the phenotypic selection coefficients on each trait included in the selection gradient analyses of *M. guttatus* and *M*. × robertsii in the reciprocal transplants experiments at each site. Significant values are in blue. The fitness measures used were fruits and stolons, respectively. Plotted values are from the models with untransformed fitness variables, and their significance is in accordance to the models with transformed variables.

#### Mimulus luteus

*M. luteus* produced significantly more fruits in IOW than in SHE (χ2 > 96.962; P < 0.001) and a similar number of stolons in both sites (χ2 = 3.329; P = 0.068). There were not differences between *M. luteus* and north *M. guttatus* in the sexual or asexual fitness overall (χ2 < 4.336; P > 0.1), but *M. luteus* produced relatively more fruits than north *M. guttatus* in SHE (χ2 = 11.018; P < 0.001). The production of stolons was higher in *M. luteus* than in north *M*. × *robertsii* overall, but it was lower in SHE (χ2 > 9.078; P < 0.003). The models of phenotypic traits showed that *M. luteus* flowered later, produced more branches and floral stems, and had lower stomata density and dry mass in SHE than in IOW (χ2 > 8.234; P < 0.01). The phenotypic traits of north *M*. × robertsii and *M. luteus* did not differ significantly, but north *M*. × *robertsii* produced relatively less branches, floral stems and flowers than *M. luteus* in SHE (negative coefficients for north *M*. × robertsii in SHE; χ2 > 9.596; P < 0.01) [Supplementary Information - Table S9]. The phenotypic selection analyses through fruit set in *M. luteus* showed significant positive linear selection in dry mass, stabilizing selection in the number of branches and dry mass, and disruptive selection in stomata density in IOW (*t* > 2.169; *P* < 0.04). In SHE, there was significant positive linear selection in the number of flowers (*t* = 2.632; *P* = 0.016) [Supplementary Information - Table S9 and Fig. S6]. The models regressing the number of stolons indicated positive linear and stabilizing selection on stomata density in IOW (*t* > 2.232; *P* < 0.035) [Supplementary Information - Table S8].

## Discussion

### Clonality, phenotypic and plasticity changes in invasive Mimulus

The reproductive systems of native *M. guttatus*, invasive *M. guttatus*, and invasive *M*. × *robertsii* showed a transition from a relatively higher investment in sexual organs (i.e. floral stems and flowers) to higher clonality (i.e. stolons). Native and invasive *M. guttatus* were similar in other phenotypic traits, suggesting that the reproductive system has been under selection in the UK, and thus supporting the important role of clonality in plant invasions (Pyšek, 1997; Song *et al*., 2013; Wang *et al*., 2017; Bock *et al*., 2018; Wang *et al*., 2019). Remarkably, annual forms of *M. guttatus* without clonal propagation do not seem to have established in the introduced range of this species. Consistent with our results, van Kleunen and Fischer (2008) found greater clonality in Scottish than in native populations of *M. guttatus*, which they related with the latitude of populations, and suggested signatures of differentiation after the species introduction at the phenotypic level. In contrast, we did not find differences in flowering time at this level as suggested by genomic analyses of selective sweeps in invasive *Mimulus* populations (Puzey and Vallejo-Marin, 2014).

Clonality has been associated with persistence at higher latitudes in *Mimulus* (Van Kleunen and Fisher, 2008) and other taxa (e.g. Dorken and Eckert, 2001). Our experiment in the controlled environment chambers allowed us to disentangle the particular drivers of this association revealing that, interestingly, warm temperatures increased clonality in both *M. guttatus* groups, but decreased clonality in *M*. × *robertsii*. Given the dependence of *M*. × *robertsii* on clonality for the long-term persistence of populations, we hypothesize that limited ability to clone in warmer environments underlies the lower abundance of *M*. × *robertsii* in the south of the UK (Hargreaves *et al*., 2014). Consistently, a previous study associated higher thermal tolerance with wider distributions in *Mimulus* (Sheth and Angert, 2014). The mechanism by which some populations of *M*. × robertsii can persist in the south of the UK (Da Re *et al*., 2020) given the reduced clonality we observe when northern populations are translocated, remains to be established.

The flowering and growth of all *Mimulus* population types were similarly affected by the temperatures and photoperiods associated to the latitudinal range of UK. In contrast, the germination of seeds was only accelerated under warm temperature treatments consistent with previous *Mimulus* work (Vickery, 1983). Warm treatments had their strongest effect on accelerating the flowering phenology of individuals, while long day treatments had their strongest effect on increasing the production of sexual organs. Warm temperatures also increased the vertical growth of plants, but not thickness nor biomass, while long photoperiods increased plant growth through all measured traits. Given the natural association of growth-promoting and growth-hindering conditions of temperature and photoperiod across latitudinal gradients, fine local microclimatic variations superimposed to large scale environmental patterns might play an important role in the performance of natural populations. In our reciprocal transplants, individuals grew bigger, flowered earlier and produced more flowers and fruits in IOW than in SHE, suggesting that, overall, the positive effects of high temperatures in the south site outperformed those of long photoperiods in the north site.

Phenotypic plasticity has been considered a distinctive trait of invasive species (Davidson *et al*., 2011) which could be also under positive selection in introduced populations (Bossdorf *et al*., 2005; Richards *et al*., 2006; but see Godoy *et al*., 2011). In our experiment, average RDPI estimates ranged 0.27-0.32 and did not differ between native and introduced populations of *M. guttatus*. This suggests a role for phenotypic plasticity as a pre-adaptation in invasive *M. guttatus* (Vickery, 1974). Native and invasive populations of *M. guttatus* showed greater phenotypic plasticity in response to temperature than to photoperiod. Given that photoperiod cycles are more constant than temperature at the local scale, this result is consistent with the hypothesis that phenotypic plasticity evolves in response to environmental variation (Via and Lande, 1985). Consisting with the classic view that clonal species rely more on phenotypic plasticity than sexual species to overcome environmental variation, our analyses indicated greater phenotypic plasticity in *M*. × *robertsii* than in *M. guttatus* (Lynch, 1984; Geng *et al*., 2007). Remarkably, the production of stolons showed similar plasticity in the two taxa, suggesting that this cannot explain their different reaction norms in the reciprocal transplants.

### *Local adaptation in introduced* Mimulus

In our reciprocal transplants experiment, we found robust patterns of local adaptation in introduced sexual populations of *M. guttatus* and asexual populations of *M*. × *robertsii*. As far as we are aware, our study is the first assessing rapid local adaptation in a strictly asexual plant species. Although there are reports of local adaptation in natural populations of other asexual multicellular organisms (e.g. Via, 1991; Ayre, 1995; Doroszuk *et al*., 2006), and in partly clonal plant populations (e.g. Lenssen *et al*., 2004), studies comparing sexual and asexual lineages are scarce and mostly based in microorganisms under laboratory conditions (e.g. Colegrave, 2002; McDonald *et al*., 2016; but see also Mariette *et al*., 2016). As a remarkable exception in plants, Lovell *et al*., (2014) compared sexual and asexual lineages of *Boechera spatifolia* but, in contrast with our study, they only found local adaptation in sexual lineages. Although we could not distinguish obvious ecotypes at each latitude for *M. guttatus* nor for *M*. × *robertsii*, and all populations were capable to survive over one season in the two extremes of the country, both species showed significant home site advantages in their respective sexual and asexual reproductive success. Parallel patterns were found in some related traits in *M. guttatus* (branches, stems, flowers) and *M*. × *robertsii* (flowers, with the opposite trend). Reproductive traits can reveal local adaptation patterns more readily than survival (Baughman *et al*., 2018), and their effects on the persistence of species are likely to act in the longer term but unequivocally. Nevertheless, the low occurrence of *M*. × *robertsii* in the south of the UK suggests that local adaptation may be more difficult to achieve in this taxon than in *M. guttatus*. Only a few populations of *M*. × *robertsii* seem to have overcome the challenges present in the south through local genotypic adaptation, which may have been facilitated by restricted dispersal opportunities (Ayre, 1995) in combination with more stressful environmental conditions (Ram and Hadany, 2002).

Our study suggest that asexual reproduction does not necessarily constrain evolution at a contemporary time scale, and this is congruent with genomic studies in other asexual plant lineages (Ferreira de Carvalho *et al*., 2016; Lovell *et al*., 2017). However, it has been also suggested that an increased heterozygosity level of hybrid polyploids in comparison with their diploid ancestors could boost their ability to adapt to different environments (Levin, 2002; Abbott *et al*., 2013; Vallejo-Marín and Hiscock, 2016; Meier et al, 2017). *M. × robertsii* differs from both parents in showing local adaptation through asexual fitness, which might propel *M × robertsii* into an independent evolutionary trajectory from its parents. Further studies are required to see if rapid local adaptation can be found also in non-hybrid asexual plants. In contrast, the fitness and phenotypic patterns of *M. luteus* were similar to those of the two other species, discarding climatic constrains on the performance of this species as explanation for its low occurrence in UK (cf. Da Re *et al*., 2020). Other environmental or ecological factors not included in our experiment, such as soil tolerance or competition with *M. × robertsii* and *M. guttatus* (Da Re *et al*., 2020), might be responsible of limiting the current distribution of *M. luteus* in the UK.

The phenotypic selection analyses showed that few and different traits were related to the sexual and/or asexual fitness of *M. guttatus* and *M*. × *robertsii* at each site in our reciprocal transplants experiment. This suggest that different mechanisms may have driven the local adaptation in each species. In *M. guttatus*, large-flowered individuals had greater sexual fitness in SHE, while shorter, heavier and late-flowering individuals had greater sexual fitness in IOW. Remarkably, flowering time is considered a principal trait under selection during species range expansions (Barrett *et al*., 2008), and in local adaptation and speciation in native *Mimulus* (e.g. Hall and Willis, 2006; Friedman and Willis, 2013). In *M*. × *robertsii*, heavier individuals had greater asexual fitness in IOW, and large-flowered individuals had greater asexual fitness in SHE. The later result contrasts with the common finding of trade-offs between sexual and asexual allocation in sexual *Mimulus* species (Sutherland and Vickery, 1988), and might be an indirect consequence of resources acquisition determined by individual quality. The selection gradients estimated for *M. luteus* were also highly different to those of *M. guttatus* and *M*. × *robertsii*. The production of fruits was positively associated with dry mass in IOW and with flower production in SHE, while the production of stolons was positively associated with stomata density in IOW. Overall, our results present partial support for a previous study which, comparing native and invasive populations of *M. guttatus*, found that introduced populations showed adaptative differentiation though selection on various traits, including large vegetative size and large floral displays and flower size (Pantoja *et al*., 2018).

The traits underlying the local adaptation of *M*. × *robertsii* in UK are yet to be fully identified, and thus populations of this species are an ideal target for further research on the mechanisms mediating rapid evolution in asexual species (see also Rushworth *et al*., 2020). Selection on clonal taxa could occur through genotypic selection in genetically diverse founding populations (clonal selection), or, perhaps, through other mechanisms including epigenetic modification (Wilschut *et al*., 2016). Although further comparisons between sexual and asexual taxa in other suitable natural systems are needed for inferences on the evolutionary rates and mechanisms of asexual taxa across plant lineages, our study provides a starting point for understanding the early evolutionary trajectory of invasive asexual plant populations.

## Acknowledgements

We thank Mathieu Quenu, Ana Montero-Castaño, Ángela Sánchez-Miranda, the V-M Lab, the University of Stirling Greenhouse staff and James Weir for their assistance during this study. We are indebted to Sally Peake for key support in Ventnor, and to Ruby Inkster from Da Gairdins i Sand and Ventnor Botanic Gardens Friend’s society for facilitating the space and performance of the reciprocal transplants. Jacinto Simba behaved very well inside the belly of VISP during the development of reciprocal transplants. Author contributions: VISP and MVM designed the study, VISP and JLS executed the experiments, VISP analysed the data and wrote the manuscript with contributions of MVM and JLS. The authors declare that there is no conflict of interest.

