## Supplementary material for "Rapid local adaptation in both sexual and asexual invasive populations of monkeyflowers (*Mimulus spp*.)"

**Table S1.** Source populations for the controlled environment chambers experiment. N indicates number of families per population. Two individuals per family (seedlings per maternal plant in *M. guttatus*; cuttings per individual in *M.* × *robertsii*), were used in the experiment. G = *M. guttatus;* R = *M. x robertsii*.

| Population | Herbarium Accession | Species | Range | Locality | Lat. | Long. | m a.s.l. | N (Families) |
|---|---|---|---|---|---|---|---|---|
| BOD | | G | Introduced | | 59.9042 | -1.3027 | 44 | 5 |
| BRA | | G | Introduced | | 52.7681 | -1.2979 | 12 | 5 |
| DBL | | G | Introduced | Dunblane, Stirlingshire | 56.1886 | -3.9661 | 64 | 5 |
| HOU | | G | Introduced | Houghton Lodge, Hampshire | 51.0970 | -1.5084 | 33 | 5 |
| TOM | | G | Introduced | | 57.2550 | -3.3678 | 318 | 5 |
| CPB | | G | Native | | 53.1710 | -131.785 | 12 | 4 |
| DAV | | G | Native | | 37.0250 | -122.2175 | 6 | 3 |
| ALA | | G | Native | | | | | |
| | V153408 | | | | 59.793 | -141.085 | | 1 |
| | V127607 | | | | 62.70 | -150.32 | | 1 |
| | V142998 | | | | 59.05 | -155.85 | | 1 |
| ORO | | G | Native | | 35.2733 | -120.8891 | 11 | 4 |
| WTB | | G | Native | | 38.4053 | -123.0961 | 35 | 4 |
| ALS | | R | Introduced | | 54.8149 | -2.4292 | 299 | 4 |
| GON | | R | Introduced | | 55.4668 | -3.7377 | 285 | 4 |
| GOO | | R | Introduced | | 57.1620 | -3.1863 | 357 | 3 |
| NEN | | R | Introduced | | 54.8061 | -2.3764 | 355 | 1 |
| WAN | | R | Introduced | | 55.3973 | -3.7804 | 405 | 3 |

**Table S2.** Temperature and photoperiod conditions of the environmental treatments implemented in the chambers experiment. Treatment codes indicate photoperiod (L, Long; S, Short) and temperature (W, Warm; C, Cold). All chambers were set to a relative humidity of 70% and a luminosity of 400 µmol·$m^{-2}$·$s^{-1}$.

| SEGMENT | | SW | | LC | | SC | | LW | |
|---|---|---|---|---|---|---|---|---|---|
| (FORTNIGTH) | | duration | temp C | duration | temp C | duration | temp C | duration | temp C |
| 1 | DAY | 12 h 54 m | 13.2 | 13 h 12 m | 7.3 | 12 h 54 m | 7.3 | 13 h 12 m | 13.2 |
| (1-15 April) | NIGTH | 11 h 6 m | 5.3 | 10 h 48 m | 2.6 | 11 h 6 m | 2.6 | 10 h 48 m | 5.3 |
| 2 | DAY | 13 h 48 m | 14.8 | 14 h 36 m | 8.6 | 13 h 48 m | 8.6 | 14 h 36 m | 14.8 |
| (16-30 April) | NIGTH | 10 h 12 m | 6.6 | 9 h 24 m | 4 | 10 h 12 m | 4 | 9 h 24 m | 6.6 |
| 3 | DAY | 14 h 42 m | 16.4 | 16 h 0 m | 9.8 | 14 h 42 m | 9.8 | 16 h 0 m | 16.4 |
| (1-15 May) | NIGTH | 9 h 18 m | 7.9 | 8 h 0 m | 5.3 | 9 h 18 m | 5.3 | 8 h 0 m | 7.9 |
| 4 | DAY | 15 h 30 m | 18.1 | 17 h 18 m | 11.2 | 15 h 30 m | 11.2 | 17 h 18 m | 18.1 |
| (16-31 May) | NIGTH | 8 h 30 m | 9.5 | 6 h 42 m | 6.7 | 8 h 30 m | 6.7 | 6 h 42 m | 9.5 |
| 5 | DAY | 16 h 12 m | 19.8 | 18 h 30 m | 12.6 | 16 h 12 m | 12.6 | 18 h 30 m | 19.8 |
| (1-15 June) | NIGTH | 7 h 48 m | 11 | 5 h 30 m | 8.1 | 7 h 48 m | 8.1 | 5 h 30 m | 11 |
| 6 | DAY | 16 h 30 m | 20.8 | 19 h 6 m | 13.1 | 16 h 30 m | 13.1 | 19 h 6 m | 20.8 |
| (16-30 June) | NIGTH | 7 h 30 m | 11.8 | 4 h 54 m | 8.8 | 7 h 30 m | 8.8 | 4 h 54 m | 11.8 |
| 7 | DAY | 16 h 24 m | 21.7 | 19 h 0 m | 13.6 | 16 h 24 m | 13.6 | 19 h 0 m | 21.7 |
| (1-15 July) | NIGTH | 7 h 36 m | 12.6 | 5 h 0 m | 9.5 | 7 h 36 m | 9.5 | 5 h 0 m | 12.6 |
| 8 | DAY | 16 h 6 m | 21.6 | 18 h 18 m | 13.6 | 16 h 6 m | 13.6 | 18 h 18 m | 21.6 |
| (16-31 July) | NIGTH | 7 h 54 m | 12.6 | 5 h 42 m | 9.5 | 7 h 54 m | 9.5 | 5 h 42 m | 12.6 |
| 9 | DAY | 15 h 24 m | 21.5 | 17 h 6 m | 13.6 | 15 h 24 m | 13.6 | 17 h 6 m | 21.5 |
| (1-15 August) | NIGTH | 8 h 36 m | 12.5 | 6 h 54 m | 9.4 | 8 h 36 m | 9.4 | 6 h 54 m | 12.5 |
| 10 | DAY | 14 h 36 m | 20.1 | 15 h 48 m | 12.4 | 14 h 36 m | 12.4 | 15 h 48 m | 20.1 |
| (16-31 August) | NIGTH | 9 h 24 m | 11.4 | 8 h 12 m | 8.4 | 9 h 24 m | 8.4 | 8 h 12 m | 11.4 |
| 11 | DAY | 13 h 36 m | 18.7 | 14 h 18 m | 11.2 | 13 h 36 m | 11.2 | 14 h 18 m | 18.7 |
| (1-15 September) | NIGTH | 10 h 24 m | 10.2 | 9 h 42 m | 7.3 | 10 h 24 m | 7.3 | 9 h 42 m | 10.2 |
| 12 | DAY | 12 h 42 m | 16.7 | 12 h 54 m | 9.9 | 12 h 42 m | 9.9 | 12 h 54 m | 16.7 |
| (16-30 September) | NIGTH | 11 h 18 m | 8.7 | 11 h 6 m | 6.1 | 11 h 18 m | 6.1 | 11 h 6 m | 8.7 |

**Table S3**. Source populations for the reciprocal transplants experiment. Species codes stand for *M. guttatus* (G) and *M.* × *robertsii* (R). The origin of populations is classified as sampled in the south or north of the British Isles.

| Population | Species | Origin | Lat. | Long. | m a.s.l. | N (individuals) |
|---|---|---|---|---|---|---|
| CRO | G | South | 50.16293 | -5.29331 | 129 | 2 |
| EAS | G | South | 50.216213 | -3.713068 | 129 | 2 |
| DAR | G | South | 50.329354 | -3.574899 | 69 | 3 |
| MOO | G | South | 50.45142 | -4.486005 | 54 | 3 |
| TCO | G | South | 50.49812 | -4.465601 | 227 | 1 |
| SOU | G | South | 50.6016 | -3.767733 | 259 | 4 |
| BOG | G | South | 50.797265 | -0.698253 | 6 | 11 |
| HUN | G | South | 50.810705 | -0.788876 | 7 | 3 |
| FUN | G | South | 50.862581 | -0.855275 | 20 | 4 |
| DEA | G | South | 50.904513 | -0.779717 | 50 | 4 |
| SIN | G | South | 50.911711 | -0.753315 | 60 | 4 |
| UPL | G | South | 50.938462 | -3.412896 | 127 | 4 |
| TOU | G | South | 51.074453 | -3.123822 | 124 | 4 |
| HOU | G | South | 51.09699 | -1.5084 | 33 | 3 |
| MOR | R | South | 50.163817 | -5.656375 | 29 | 5 |
| DRI | R | South | 51.148021 | -3.808392 | 384 | 6 |
| MAR | G | North | 57.572341 | -4.427486 | 3 | 5 |
| GAR | G | North | 57.615064 | -4.673473 | 75 | 4 |
| DAL | G | North | 57.682614 | -4.265258 | 6 | 4 |
| BLA | G | North | 58.48755 | -5.10636 | 44 | 4 |
| BKN | G | North | 58.5759 | -4.76774 | 8 | 4 |
| BOD | G | North | 59.90418 | -1.30274 | 55 | 11 |
| NIN | G | North | 59.97777 | -1.30036 | 87 | 5 |
| WEI | G | North | 60.254393 | -1.289859 | 6 | 4 |
| MUK | G | North | 60.34808 | -1.41373 | 8 | 4 |
| HAM | G | North | 60.5034 | -1.09931 | 4 | 4 |
| NORG | G | North | 60.808743 | -0.807753 | 13 | 5 |
| GJO | G | North | 62.32533 | -6.94162 | 3 | 2 |
| CLS | R | North | 58.215153 | -5.33411 | 46 | 7 |
| GIO | R | North | 58.33754 | -6.20187 | 38 | 2 |
| POL | R | North | 58.483497 | -5.099521 | 20 | 11 |
| EOR | R | North | 58.49937 | -6.26996 | 8 | 3 |
| STR | R | North | 58.9692 | -3.28341 | 9 | 3 |
| TOR | R | North | 58.9957 | -3.18338 | 26 | 2 |
| EVI | R | North | 59.11226 | -3.10809 | 37 | 1 |
| NOR | R | North | 60.808743 | -0.807753 | 13 | 7 |
| COL | L | North | 55.6550 | -2.2401 | 9 | 25 |

**Table S4.** Summary of results of the GLMMs modelling the variation in all traits measured in the controlled environmental chambers as a function of temperature, photoperiod, population type (where appropriate) and their interactions.

| Trait | Fixed factor | | All populations Estimate (SE) | $\chi^2$ | Native *M. guttatus* Estimate (SE) | $\chi^2$ | Invasive *M. guttatus* Estimate (SE) | $\chi^2$ | M. × *robertsii* Estimate (SE) | $\chi^2$ |
|---|---|---|---|---|---|---|---|---|---|---|
| **Germination day** | Intercept | | 13.475 (0.498) | | 13.2 (0.353) | | 13.41 (0.65) | | | |
| | **T** | **(Warm)** | -6.087 (0.25) | **590.496 ***** | -6.2 (0.408) | **231.463 ***** | -6.02 (0.314) | **367.683 ***** | | |
| | **P** | **(Short)** | 0.237 (0.25) | 0.899 | 0.133 (0.408) | 0.107 | 0.3 (0.314) | 0.913 | | |
| | **O** | **(US)** | -0.366 (0.707) | 0.268 | | | | | | |
| **Flower day** | Intercept | | 91.183 (2.248) | | 87.645 (1.589) | | 90.637 (1.906) | | 81.767 (4.25) | |
| | **T** | **(Warm)** | -22.014 (1.279) | **296.237 ***** | -24.022 (1.504) | **255.145 ***** | -22.294 (1.111) | **402.322 ***** | -17.892 (4.976) | **12.93 ***** |
| | **P** | **(Short)** | 8.52 (1.297) | **43.156 ***** | 9.949 (1.504) | **43.763 ***** | 10.018 (1.114) | **80.876 ***** | 0.443 (5.221) | 0.007 |
| | **O** | **(US)** | -3.838 (2.986) | 8.298 * | | | | | | |
| | | **(rob)** | -9.103 (3.16) | | | | | | | |
| **Flowered** | Intercept | | 1.031 (0.052) | | 1.001 (0.032) | | 1.00 (0.044) | | 0.972 (0.099) | |
| | **T** | **(Warm)** | -0.015 (0.028) | 0.289 | -0.013 (0.03) | 0.194 | -0.02 (0.033) | 0.356 | -0.01 (0.079) | 0.016 |
| | **P** | **(Short)** | -0.167 (0.028) | **33.538 ***** | -0.042 (0.03) | 2.055 | -0.1 (0.033) | **8.911 **** | -0.423 (0.079) | **28.559 ***** |
| | **O** | **(US)** | 0.032 (0.07) | 10.339 ** | | | | | | |
| | | **(rob)** | -0.186 (0.071) | | | | | | | |
| **Corolla** | Intercept | | 38.972 (1.42) | | 39.274 (1.258) | | 38.888 (2.016) | | 35.9 (1.09) | |
| | **T** | **(Warm)** | -2.637 (0.496) | **28.25 ***** | -2.693 (1.05) | 6.571 | -3.298 (0.589) | **31.361 ***** | -1.19 (1.015) | 1.375 |
| | **P** | **(Short)** | -1.254 (0.502) | 6.233 * | -2.684 (1.05) | 6.528 | -0.403 (0.59) | 0.467 | -0.899 (1.061) | 0.717 |
| | **O** | **(US)** | -0.479 (1.963) | 1.43 | | | | | | |
| | | **(rob)** | -2.3 (2.01) | | | | | | | |

| | | | | | | | | | | |
|---|---|---|---|---|---|---|---|---|---|---|
| **Flowers** | Intercept | | 3.539 (0.118) | | 3.675 (0.13) | | 3.54 (0.079) | | 2.037 (0.178) | |
| | **T** | **(Warm)** | 0.128 (0.046) | 7.713 ** | 0.315 (0.049) | **41.564 *** ** | 0.129 (0.046) | 7.712 | 0.199 (0.102) | 3.797 |
| | **P** | **(Short)** | -0.87 (0.063) | **186.062 *** ** | -0.496 (0.061) | **67.063 *** ** | -0.87 (0.064) | **185.731 *** ** | -0.654 (0.117) | **31.037 *** ** |
| | **T : P** | **(Warm:Short)** | 0.313 (0.082) | **14.412 *** ** | 0.348 (0.077) | **20.612 *** ** | 0.313 (0.083) | **14.343 *** ** | | |
| | **O** | **(US)** | 0.136 (0.168) | **81.676 *** ** | | | | | | |
| | | **(rob)** | -1.462 (0.189) | | | | | | | |
| | **O : T** | **(US:Warm)** | 0.186 (0.067) | 7.883 * | | | | | | |
| | | **(rob:Warm)** | 0.027 (0.127) | | | | | | | |
| | **O : P** | **(US:Short)** | 0.375 (0.087) | **18.305 *** ** | | | | | | |
| | | **(rob:Short)** | 0.141 (0.184) | | | | | | | |
| | **O : T : P** | **(US:W:S)** | 0.033 (0.112) | 0.617 | | | | | | |
| | | **(rob:W:S)** | -0.154 (0.245) | | | | | | | |
| **Stems** | Intercept | | 1.436 (0.095) | | 1.786 (0.115) | | 1.62 (0.092) | | 0.782 (0.179) | |
| | **T** | **(Warm)** | -0.089 (0.072) | 1.532 | 0.072 (0.104) | 0.474 | -0.287 (0.114) | 6.323 | -0.061 (0.211) | 0.084 |
| | **P** | **(Short)** | -0.571 (0.076) | **56.545 *** ** | -0.373 (0.106) | **12.357 *** ** | -0.847 (0.125) | **46.294 *** ** | -0.476 (0.238) | 4.002 |
| | **O** | **(US)** | 0.506 (0.118) | **57.637 *** ** | | | | | | |
| | | **(rob)** | -0.612 (0.151) | | | | | | | |
| **Height** | Intercept | | 29.5 (2.383) | | 33.459 (2.017) | | 31.445 (2.866) | | 20.969 (2.09) | |
| | **T** | **(Warm)** | 23.48 (2.132) | **121.317 *** ** | 15.683 (1.888) | **68.976 *** ** | 19.59 (1.605) | **148.986 *** ** | 7.217 (1.775) | **16.535 *** ** |
| | **P** | **(Short)** | -2.86 (2.132) | 1.8 | -4.905 (1.888) | 6.747 | -6.75 (1.605) | **17.688 *** ** | -10.531 (1.782) | **34.93 *** ** |
| | **T : P** | **(Warm:Short)** | -7.78 (3.015) | 6.66 ** | | | | | | |
| | **O** | **(US)** | 3.38 (3.501) | **13.928 *** ** | | | | | | |
| | | **(rob)** | -10.076 (3.614) | | | | | | | |
| | **O : T** | **(US:Warm)** | -6.839 (3.325) | **14.855 *** ** | | | | | | |
| | | **(rob:Warm)** | -13.247 (3.481) | | | | | | | |
| | **O : P** | **(US:Short)** | -1.088 (3.325) | 1.847 | | | | | | |
| | | **(rob:Short)** | -4.677 (3.486) | | | | | | | |
| | **O : T : P** | **(US:W:S)** | 5.865 (4.704) | 1.575 | | | | | | |
| | | **(rob:W:S)** | 1.747 (4.923) | | | | | | | |

| | | | | | | | | | | |
|---|---|---|---|---|---|---|---|---|---|---|
| **Branches** | Intercept | | 0.46 (0.196) | | 0.081 (0.32) | | 0.414 (0.138) | | -0.429 (0.323) | |
| | **T** | **(Warm)** | 0.322 (0.107) | **9.094 **** | 0.339 (0.182) | 3.487 | 0.373 (0.142) | 6.932 | -0.154 (0.392) | 0.154 |
| | **P** | **(Short)** | 0.128 (0.105) | 1.474 | 0.203 (0.18) | 1.271 | 0.195 (0.14) | 1.936 | -0.799 (0.425) | 3.534 |
| | **O** | **(US)** | -0.275 (0.262) | **24.271 ***** | | | | | | |
| | | **(rob)** | -1.521 (0.313) | | | | | | | |
| **Internode length** | Intercept | | 14.128 (0.971) | | 13.671 (1.021) | | 14.492 (1.004) | | 10.283 (1.787) | |
| | **T** | **(Warm)** | 2.197 (0.727) | **9.134 **** | 0.498 (1.19) | 0.175 | 3.322 (0.912) | **13.273 ***** | 2.315 (1.773) | 1.704 |
| | **P** | **(Short)** | -3.833 (0.727) | **27.761 ***** | -4.081 (1.19) | **11.753 ***** | -5.686 (0.912) | **38.884 ***** | -0.402 (1.778) | 0.051 |
| | **O** | **(US)** | -1.503 (1.224) | 2.988 | | | | | | |
| | | **(rob)** | -2.07 (1.269) | | | | | | | |
| **Internode diameter** | Intercept | | 7.234 (0.212) | | 6.808 (0.33) | | 7.382 (0.203) | | 3.908 (0.26) | |
| | **T** | **(Warm)** | -0.638 (0.145) | **19.311 ***** | -0.786 (0.294) | 7.1424 | -0.853 (0.221) | **14.869 ***** | -0.103 (0.214) | 0.232 |
| | **P** | **(Short)** | -0.654 (0.145) | **20.268 ***** | -0.798 (0.294) | 7.357 | -0.733 (0.221) | **10.98 ***** | -0.354 (0.215) | 2.712 |
| | **O** | **(US)** | -0.562 (0.272) | **116.882 ***** | | | | | | |
| | | **(rob)** | -2.934 (0.281) | | | | | | | |
| **Drymass** | Intercept | | 5.199 (0.528) | | 4.28 (0.299) | | 5.36 (0.214) | | 5.185 (0.959) | |
| | **T** | **(Warm)** | -0.545 (0.173) | **9.891 **** | -0.122 (0.206) | 0.349 | -0.7 (0.163) | **18.521 ***** | -0.787 (0.549) | 2.054 |
| | **P** | **(Short)** | -0.606 (0.173) | **12.206 ***** | -0.373 (0.206) | 3.259 | -0.774 (0.163) | **22.67 ***** | -0.609 (0.55) | 1.226 |
| | **O** | **(US)** | -0.596 (0.732) | 0.72 | | | | | | |
| | | **(rob)** | -0.137 (0.739) | | | | | | | |
| **Stolons** | Intercept | | 1.744 (0.209) | | 1.26 (0.332) | | 1.75 (0.112) | | 2.616 (0.126) | |
| | **T** | **(Warm)** | 0.319 (0.108) | **8.67 **** | 0.872 (0.138) | **39.749 ***** | 0.32 (0.109) | 8.6635 | -0.234 (0.073) | **10.222** |
| | **P** | **(Short)** | 0.231 (0.11) | 4.369 * | 0.332 (0.154) | 4.644 | 0.231 (0.111) | 4.3664 | -0.005 (0.073) | 0.005 |
| | **T : P** | **(Warm:Short)** | -0.369 (0.151) | 5.972 * | -0.756 (0.196) | **14.909 ***** | -0.37 (0.151) | 5.9679 | | |
| | **O** | **(US)** | -0.468 (0.308) | **17.974 ***** | | | | | | |

| | | | |
|---|---|---|---|
| | **(rob)** | 0.801 (0.294) | |
| **O : T** | **(US:Warm)** | 0.549 (0.175) | **32.035 *** ** |
| | **(rob:Warm)** | -0.418 (0.148) | |
| **O : P** | **(US:Short)** | 0.096 (0.189) | 1.452 |
| | **(rob:Short)** | -0.111 (0.147) | |
| **O : T : P** | **(US:W:S)** | -0.379 (0.247) | |
| | **(rob:W:S)** | 0.092 (0.21) | 3.881 |

**Table note:** Fixed effect abreviations: Origin (O); Temperature (T); Photoperiod (P). $\chi$2 values and their associated *P*-values from type-III Wald $\chi$2 tests (or type-II Wald $\chi$2 tests, when all interaction terms were unsignificant and removed) are provided. * $P<0.05$; ** $P<0.01$; *** $P<0.001$. Significant effects after Bonferroni correction of *P*-values are indicated in bold.

**Table S5.** Mean value, standard deviation and sample size for each trait and group defined by a significant fixed effect in the GLMMs modelling the phenotypic data for all population types in the controlled environmental chambers.

| Trait | Fixed effect | Group | mean ± sd (N) |
|---|---|---|---|
| **Germination day** | **T** | Cold | 13.45 ± 2.349 (80) |
| | | Warm | 7.363 ± 1.343 (80) |
| **Flower day** | **T** | Cold | 91.461 ± 11.811 (103) |
| | | Warm | 69.622 ± 11.311 (107) |
| | **P** | Long | 76.662 ± 14.635 (114) |
| | | Short | 84.693 ± 16.305 (96) |
| **Flowered** | **P** | Long | 0.987 ± 0.08 (115) |
| | | Short | 0.819 ± 0.351 (116) |
| **Corolla** | **T** | Cold | 38 ± 5.289 (106) |
| | | Warm | 35.275 ± 4.61 (106) |
| **Flowers** | **O** | *M. guttatus* invasive | 28.332 ± 16.368 (98) b |
| | | *M. guttatus* native | 43.333 ± 22.34 (69) c |
| | | *M. x robertsii* | 7.929 ± 6.009 (49) a |
| | **P** | Long | 33.622 ± 23.02 (115) |
| | | Short | 22.658 ± 17.523 (101) |
| | **O:P** | *M. guttatus* invasive : Long | 37.33 ± 16.892 (50) de |
| | | *M. guttatus* invasive : Short | 18.958 ± 8.923 (48) c |
| | | *M. guttatus* native : Long | 49 ± 23.446 (35) e |
| | | *M. guttatus* native : Short | 37.5 ± 19.821 (34) d |
| | | *M. x robertsii* : Long | 9.5 ± 6.745 (30) b |
| | | *M. x robertsii* : Short | 5.447 ± 3.515 (19) a |
| | **T:P** | Cold : Long | 30.034 ± 18.115 (58) c |
| | | Cold : Short | 16.48 ± 11.116 (50) a |
| | | Warm : Long | 37.272 ± 26.793 (57) d |
| | | Warm : Short | 28.716 ± 20.43 (51) b |
| **Stems** | **O** | *M. guttatus invasive* | 3.179 ± 2.179 (98) b |
| | | *M. guttatus native* | 5.239 ± 2.182 (69) c |
| | | *M. x robertsii* | 1.816 ± 1.121 (49) a |
| **Height** | **O** | *M. guttatus invasive* | 37.865 ± 14.325 (100) b |
| | | *M. guttatus native* | 39.186 ± 11.725 (70) b |
| | | *M. x robertsii* | 19.475 ± 9.861 (60) a |
| | **T** | Cold | 25.904 ± 9.904 (115) |

| | | | |
|---|---|---|---|
| | | Warm | 41.035 ± 15.436 (115) |
| | **P** | Long | 4.396 ± 2.486 (115) |
| | | Short | 2.54 ± 1.767 (101) |
| | **O:T** | *M. guttatus* invasive : Cold | 28.07 ± 7.465 (50) b |
| | | *M. guttatus* invasive : Warm | 47.66 ± 12.769 (50) c |
| | | *M. guttatus* native : Cold | 31.414 ± 7.832 (35) b |
| | | *M. guttatus* native : Warm | 46.957 ± 9.66 (35) c |
| | | *M. x robertsii* : Cold | 15.867 ± 8.398 (30) a |
| | | *M. x robertsii* : Warm | 23.083 ± 10.017 (30) b |
| **Branches** | **O** | *M. guttatus* invasive | 2.01 ± 1.409 (100) b |
| | | *M. guttatus* native | 1.779 ± 1.634 (70) b |
| | | *M. x robertsii* | 0.458 ± 0.825 (60) a |
| | **T** | Cold | 1.27 ± 1.372 (115) |
| | | Warm | 1.8 ± 1.585 (115) |
| **Internode** | **T** | Cold | 11.259 ± 6.199 (115) |
| | | Warm | 13.423 ± 5.949 (115) |
| | **P** | Long | 14.233 ± 6.136 (115) |
| | | Short | 10.45 ± 5.595 (115) |
| **Diameter** | **T** | Cold | 5.976 ± 1.791 (115) |
| | | Warm | 5.332 ± 1.651 (115) |
| | **P** | Long | 5.982 ± 1.741 (115) |
| | | Short | 5.325 ± 1.701 (115) |
| | **O** | *M. guttatus* invasive | 6.588 ± 1.256 (100) b |
| | | *M. guttatus* native | 6.068 ± 1.443 (70) b |
| | | *M. x robertsii* | 3.613 ± 0.931 (60) a |
| **Drymass** | **T** | Cold | 4.704 ± 1.619 (115) |
| | | Warm | 4.142 ± 1.797 (116) |
| | **P** | Long | 4.737 ± 1.701 (115) |
| | | Short | 4.11 ± 1.708 (116) |
| **Stolons** | **O** | *M. guttatus* invasive | 7.025 ± 3.018 (100) ab |
| | | *M. guttatus* native | 6.407 ± 7.506 (70) a |
| | | *M. x robertsii* | 12.558 ± 5.899 (60) b |
| | **T** | Cold | 7.996 ± 6.203 (115) |
| | | Warm | 8.565 ± 5.878 (115) |
| | **O:T** | *M. guttatus* invasive : Cold | 6.56 ± 2.697 (50) abc |
| | | *M. guttatus* invasive : Warm | 7.49 ± 3.269 (50) abc |

| | | |
|---|---|---|
| | *M. guttatus* native : Cold | 4.871 ± 6.543 (35) a |
| | *M. guttatus* native: Warm | 7.943 ± 8.165 (35) bc |
| | *M. x robertsii* : Cold | 14.033 ± 5.975 (30) c |
| | *M. x robertsii* : Warm | 11.083 ± 5.531 (30) b |

**Table note:** Fixed effect abreviations: Origin (O); Temperature (T); Photoperiod (P). When more than two groups are defined by the significant fixed term, significantly different groups according to *post hoc* tests are indicated with different letters.

**Table S6.** Results of the GLMMs analysing the RDPI indexes of phenotypic plasticity as a function of the population type for all traits measured in the controlled environmental chambers.

| **(a) RDPIt** | $\chi^2$ | df | *P* | *M. guttatus* native | *M. guttatus* invasive | *M.* x *robertsii* |
|---|---|---|---|---|---|---|
| Branches | 8.679 | 2 | 0.013 * | 0.636 ± 0.304 (18) | 0.469 ± 0.297 (25) | 0.821 ± 0.244 (13) |
| Flower day | 1.015 | 2 | 0.602 | 0.26 ± 0.042 (18) | 0.23 ± 0.055 (25) | 0.239 ± 0.185 (13) |
| Germination day | 2.106 | 1 | 0.147 | 0.466 ± 0.035 (15) | 0.441 ± 0.064 (25) | -- |
| Stems | 5.552 | 2 | 0.062 | 0.227 ± 0.171 (18) | 0.395 ± 0.222 (25) | 0.214 ± 0.202 (15) |
| Corolla width | 0.15 | 2 | 0.928 | 0.093 ± 0.056 (18) | 0.091 ± 0.054 (25) | 0.089 ± 0.054 (13) |
| Height | 5.064 | 2 | 0.08 | 0.365 ± 0.117 (18) | 0.404 ± 0.098 (25) | 0.307 ± 0.202 (16) |
| Diameter | 0.872 | 2 | 0.647 | 0.169 ± 0.143 (18) | 0.139 ± 0.104 (25) | 0.165 ± 0.114 (16) |
| Flowers | 5.02 | 2 | 0.081 | 0.428 ± 0.156 (18) | 0.296 ± 0.19 (25) | 0.431 ± 0.24 (15) |
| Internode | 3.256 | 2 | 0.196 | 0.32 ± 0.109 (18) | 0.273 ± 0.138 (25) | 0.353 ± 0.183 (16) |
| Drymass | 3.468 | 2 | 0.177 | 0.159 ± 0.118 (18) | 0.175 ± 0.132 (25) | 0.247 ± 0.201 (16) |
| Stolons | 4.321 | 2 | 0.115 | 0.426 ± 0.198 (18) | 0.273 ± 0.178 (25) | 0.317 ± 0.242 (16) |

| **(b) RDPIp** | Chisq | df | *P* | *M. guttatus* snative | *M. guttatus* invasive | *M.* x *robertsii* |
|---|---|---|---|---|---|---|
| Branches | 18.272 | 2 | < 0.001 *** | **0.46 ± 0.355 (18) a** | **0.341 ± 0.188 (25) a** | **0.797 ± 0.366 (13) b** |
| Flower day | 23.334 | 2 | < 0.001 *** | **0.128 ± 0.054 (18) a** | **0.109 ± 0.061 (25) a** | **0.237 ± 0.123 (10) b** |
| Germination day | 0.007 | 1 | 0.933 | 0.055 ± 0.054 (15) | 0.056 ± 0.045 (25) | -- |
| Stems | 16.526 | 2 | < 0.001 *** | **0.335 ± 0.174 (18) a** | **0.575 ± 0.204 (25) b** | **0.445 ± 0.219 (11) ab** |
| Corolla width | 2.77 | 2 | 0.25 | 0.085 ± 0.064 (18) | 0.068 ± 0.047 (25) | 0.055 ± 0.035 (11) |
| Height | 15.837 | 2 | < 0.001 *** | **0.169 ± 0.135 (18) a** | **0.199 ± 0.159 (25) a** | **0.426 ± 0.199 (16) b** |
| Diameter | 0.214 | 2 | 0.899 | 0.158 ± 0.116 (18) | 0.145 ± 0.101 (25) | 0.141 ± 0.112 (16) |
| Flowers | 6.201 | 2 | 0.045 * | 0.315 ± 0.18 (18) | 0.471 ± 0.199 (25) | 0.484 ± 0.216 (11) |
| Internode | 0.405 | 2 | 0.817 | 0.31 ± 0.137 (18) | 0.343 ± 0.176 (25) | 0.317 ± 0.22 (16) |
| Drymass | 7.342 | 2 | 0.025 * | 0.162 ± 0.102 (18) | 0.164 ± 0.09 (25) | 0.283 ± 0.215 (16) |
| Stolons | 1.342 | 2 | 0.511 | 0.303 ± 0.19 (18) | 0.289 ± 0.139 (25) | 0.243 ± 0.16 (16) |

| **(c) RDPItp** | Chisq | df | *P* | *M. guttatus* native | *M. guttatus* invasive | *M.* x *robertsii* |
|---|---|---|---|---|---|---|
| Branches | 1.453 | 2 | 0.484 | 0.527 ± 0.209 (13) | 0.543 ± 0.176 (24) | 0.685 ± 0.085 (13) |
| Flower day | 5.123 | 2 | 0.077 | 0.209 ± 0.028 (16) | 0.198 ± 0.042 (21) | 0.247 ± 0.095 (6) |
| Germination day | 0.893 | 1 | 0.345 | 0.332 ± 0.037 (15) | 0.319 ± 0.044 (25) | -- |
| Stems | 7.756 | 2 | 0.021 * | 0.342 ± 0.153 (16) | 0.498 ± 0.092 (23) | 0.364 ± 0.211 (7) |
| Corolla width | 2.069 | 2 | 0.355 | 0.105 ± 0.038 (16) | 0.095 ± 0.029 (22) | 0.078 ± 0.056 (6) |
| Height | 4.747 | 2 | 0.093 | 0.293 ± 0.099 (17) | 0.34 ± 0.07 (25) | 0.389 ± 0.172 (12) |
| Diameter | 1.203 | 2 | 0.548 | 0.21 ± 0.097 (17) | 0.18 ± 0.074 (25) | 0.202 ± 0.104 (12) |
| Flowers | 0.646 | 2 | 0.724 | 0.417 ± 0.097 (16) | 0.431 ± 0.11 (23) | 0.456 ± 0.14 (7) |
| Internode | 0.169 | 2 | 0.919 | 0.347 ± 0.102 (17) | 0.349 ± 0.104 (25) | 0.371 ± 0.184 (12) |
| Drymass | 13.655 | 2 | 0.001 ** | **0.207 ± 0.068 (17) a** | **0.215 ± 0.091 (25) a** | **0.319 ± 0.126 (13) b** |
| Stolons | 5.034 | 2 | 0.081 | 0.43 ± 0.114 (17) | 0.34 ± 0.131 (25) | 0.327 ± 0.172 (12) |

**Table note:** Mean ± s.d. (N) is given for each trait and group, and significant differences in the *post hoc* tests are labelled with different letters. * P<0.05; ** P<0.01; *** P<0.001.

**Table S7**. Results of the GLMMs modelling the effects of initial weigth ($W_0$), population origin, experimental site and their interaction in 10 phenotypic and phenological traits recorded in a reciprocal transplant experiment with introduced *Mimulus guttatus* and *M.* × *robertsii* populations.

| | | ***M. guttatus*** | | ***M.* × *robertsii*** | |
|---|---|---|---|---|---|
| **Trait** | **Fixed factor** | **Estimate (SE)** | $\chi^2$ | **Estimate (SE)** | $\chi^2$ |
| **Flowering day** | Intercept | 49.206 (2.227) | 488.243 *** | 37.919 (2.272) | 278.455 *** |
| | $\mathbf{W_0}$ | -0.053 (0.021) | 6.522 * | -0.007 (0.006) | 1.202 |
| | **Origin** (South) | -0.71 (1.691) | 0.176 | 5.622 (4.715) | 1.422 |
| | **Site** (SHE) | 24.121 (1.017) | **562.793 ***** | 25.529 (1.354) | **355.649 ***** |
| | **S:O** (SHE:South) | -0.66 (1.271) | 0.269 | -5.415 (2.739) | 3.908 * |
| **Flowering node** | Intercept | 5.882 (0.433) | 184.892 *** | 4.536 (0.265) | 292.421 *** |
| | $\mathbf{W_0}$ | 0.002 (0.003) | 0.219 | 0 (0.001) | 0.212 |
| | **Origin** (South) | 0.031 (0.415) | 0.006 | 0.256 (0.53) | 0.234 |
| | **Site** (SHE) | -0.103 (0.168) | 0.376 | 0.132 (0.172) | 0.59 |
| | **S:O** (SHE:South) | -0.053 (0.208) | 0.064 | -0.303 (0.344) | 0.774 |
| **Branches** | Intercept | 2.394 (0.116) | 422.442 *** | 2.158 (0.079) | 746.99 *** |
| | $\mathbf{W_0}$ | -0.002 (0.001) | 5.303 * | 0.001 (0) | 4.544 * |
| | **Origin** (South) | 0.142 (0.093) | 2.307 | 0 (0.15) | 0 |
| | **Site** (SHE) | 0.011 (0.051) | 0.044 | -0.073 (0.057) | 1.682 |
| | **S:O** (SHE:South) | -0.197 (0.062) | **10.14 **** | 0.207 (0.11) | 3.514 |
| **Stems** | Intercept | 2.844 (0.111) | 655.131 *** | 2.635 (0.081) | 1053.154 *** |
| | $\mathbf{W_0}$ | -0.002 (0.001) | 4.087 * | 0.001 (0) | **12.247 ***** |
| | **Origin** (South) | 0.095 (0.115) | 0.679 | 0.014 (0.157) | 0.008 |
| | **Site** (SHE) | -0.046 (0.04) | 1.353 | -0.13 (0.045) | **8.4 **** |
| | **S:O** (SHE:South) | -0.153 (0.048) | **9.933 **** | 0.146 (0.087) | 2.822 |
| **Flowers** | Intercept | 4.915 (0.105) | 2179.201 *** | 5.093 (0.076) | 4455.005 *** |
| | $\mathbf{W_0}$ | -0.001 (0) | **11.943 ***** | 0 (0) | **42.282 ***** |
| | **Origin** (South) | 0.196 (0.142) | 1.909 | -0.017 (0.159) | 0.012 |
| | **Site** (SHE) | -0.181 (0.015) | **147.487 ***** | -0.146 (0.014) | **116.622 ***** |
| | **S:O** (SHE:South) | -0.101 (0.018) | **32.664 ***** | 0.105 (0.027) | **15.11 ***** |
| **Corolla** | Intercept | 399.725 (18.206) | 482.072 *** | 422.987 (15.085) | 786.301 *** |
| | $\mathbf{W_0}$ | -0.031 (0.165) | 0.036 | -0.075 (0.042) | 3.141 |
| | **Origin** (South) | -38.025 (14.839) | 6.566 * † | -44.584 (30.787) | 2.097 |
| | **Site** (SHE) | -14.039 (8.091) | 3.011 | 13.015 (9.624) | 1.829 |
| | **S:O** (SHE:South) | -4.286 (10.07) | 0.181 | -41.657 (19.368) | 4.626 * |

| | | | | | |
|---|---|---|---|---|---|
| **Stomata** | Intercept | 18.137 (1.03) | 310.282 *** | 14.336 (1.095) | 171.362 *** |
| | **$W_0$** | -0.034 (0.01) | **10.598 **** | 0.001 (0.002) | 0.285 |
| | **Origin** (South) | -0.459 (0.656) | 0.489 | -1.779 (2.334) | 0.581 |
| | **Site** (SHE) | -2.555 (0.516) | **24.524 ***** | -0.195 (0.525) | 0.138 |
| | **S:O** (SHE:South) | -0.171 (0.651) | 0.069 | 0.25 (1.059) | 0.056 |
| **Height** | Intercept | 19.816 (2.646) | 56.103 *** | 22.081 (1.348) | 268.363 *** |
| | **$W_0$** | 0.035 (0.024) | 2.142 | 0.02 (0.007) | **9.728 **** |
| | **Origin** (South) | 2.904 (2.153) | 1.82 | 0.424 (2.358) | 0.032 |
| | **Site** (SHE) | -3.204 (1.167) | 7.544 ** | -0.402 (1.411) | 0.081 |
| | **S:O** (SHE:South) | 2.763 (1.453) | 3.616 | 2.26 (2.852) | 0.628 |
| **Cover** | Intercept | 906.806 (74.199) | 149.36 *** | 640.669 (52.4) | 149.488 *** |
| | **$W_0$** | -2.091 (0.667) | **9.847 **** | 0.41 (0.172) | 5.688 * |
| | **Origin** (South) | 8.063 (62.149) | 0.017 | 48.324 (104.377) | 0.214 |
| | **Site** (SHE) | -267.054 (32.495) | **67.541 ***** | -59.204 (38.272) | 2.393 |
| | **S:O** (SHE:South) | -14.122 (40.777) | 0.12 | -46.233 (77.581) | 0.355 |
| **Dry mass** | Intercept | 37.479 (4.929) | 57.814 *** | 35.644 (3.015) | 139.769 *** |
| | **$W_0$** | 0.013 (0.046) | 0.075 | 0.014 (0.009) | 2.567 |
| | **Origin** (South) | -0.511 (3.793) | 0.018 | 1.745 (6.092) | 0.082 |
| | **Site** (SHE) | -15.924 (2.25) | **50.089 ***** | -9.85 (2.029) | **23.579 ***** |
| | **S:O** (SHE:South) | 0.676 (2.804) | 0.058 | -2.334 (4.106) | 0.323 |

**Table note:** Models were built on each individual species dataset. Maximal model Estimates and Standard Errors for each fixed factor and interaction are provided jointly with results of the type-III Wald $\chi 2$ tests. $\chi 2$ values and indications of their associated *P*-values are provided. * P < 0.05; ** P < 0.01; *** P < 0.001. Significant effects after Bonferroni correction of *P*-values are indicated in bold. $\chi 2$ degrees of freedom=1. † Indicates significant main effects after type-II Wald $\chi 2$ tests on GLMMs excluding non-significant interactions (only significant effects after Bonferroni correction are shown).

**Table S8.** Results of the regression models of phenotypic selection on 10 traits measured in *Mimulus guttatus*, *M.* × *robertsii* and *M. luteus* populations included in the reciprocal transplants experiment.

| **(a) *M. guttatus*** | **Fruits** | | | | **Stolons** | | | |
|---|---|---|---|---|---|---|---|---|
| | **IOW** | | **SHE** | | **IOW** | | **SHE** | |
| | **Estimate (SE)** | ***t*** | **Estimate (SE)** | ***t*** | **Estimate (SE)** | ***t*** | **Estimate (SE)** | ***t*** |
| Intercept | 0.868 (0.029) | 29.955 *** | 0.82 (0.039) | 20.925 *** | 0.859 (0.037) | 22.951 *** | 0.849 (0.043) | 19.857 *** |
| Height | -0.392 (0.16) | **-2.453 *** | -0.086 (0.207) | -0.416 | -0.084 (0.207) | -0.405 | 0.142 (0.233) | 0.611 |
| Stomata | -0.024 (0.189) | -0.127 | -0.098 (0.256) | -0.385 | 0.104 (0.244) | 0.428 | -0.197 (0.287) | -0.687 |
| Flowering day | 0.57 (0.217) | **2.631 **** | 0.534 (0.491) | 1.088 | -0.029 (0.277) | -0.106 | -0.235 (0.451) | -0.521 |
| Flowering node | 0.293 (0.199) | 1.471 | 0.003 (0.261) | 0.011 | 0.302 (0.259) | 1.168 | 0.311 (0.24) | 1.297 |
| Corolla | 0.187 (0.292) | 0.64 | 0.671 (0.291) | **2.305 *** | -0.326 (0.379) | -0.86 | -0.201 (0.325) | -0.619 |
| Plant cover | -0.056 (0.155) | -0.363 | 0.173 (0.242) | 0.716 | 0.029 (0.199) | 0.147 | 0.362 (0.271) | 1.337 |
| Branches | -0.175 (0.143) | -1.223 | 0.079 (0.235) | 0.334 | -0.223 (0.186) | -1.198 | -0.618 (0.249) | **-2.482 *** |
| Floral stems | 0.27 (0.152) | 1.781 | -0.156 (0.209) | -0.745 | 0.314 (0.196) | 1.603 | 0.398 (0.232) | 1.718 |
| Flowers | -0.134 (0.126) | -1.062 | 0.135 (0.149) | 0.902 | -0.277 (0.163) | -1.697 | -0.217 (0.165) | -1.317 |
| Dry mass | 0.412 (0.107) | **3.864 ***** | 0.099 (0.191) | 0.519 | 0.142 (0.137) | 1.033 | -0.477 (0.216) | **-2.203 *** |
| Height^2 | 0.743 (0.309) | **2.407 *** | -0.015 (0.385) | -0.039 | 0.121 (0.401) | 0.301 | -0.159 (0.439) | -0.363 |
| Stomata^2 | 0.019 (0.38) | 0.051 | 0.199 (0.52) | 0.383 | -0.118 (0.491) | -0.241 | 0.461 (0.578) | 0.797 |
| Flowering day^2 | -1.328 (0.447) | **-2.97 **** | -1.296 (1.028) | -1.262 | 0.106 (0.567) | 0.187 | 0.634 (0.912) | 0.695 |
| Flowering node^2 | -0.591 (0.394) | -1.501 | -0.196 (0.506) | -0.388 | -0.613 (0.511) | -1.201 | -0.552 (0.469) | -1.178 |
| Corolla^2 | -0.333 (0.589) | -0.566 | -1.101 (0.576) | -1.912 | 0.651 (0.764) | 0.852 | 0.529 (0.65) | 0.814 |
| Plant cover^2 | 0.248 (0.284) | 0.874 | 0.048 (0.406) | 0.118 | 0.057 (0.366) | 0.155 | -0.459 (0.454) | -1.011 |
| Branches^2 | 0.361 (0.287) | 1.258 | -0.268 (0.54) | -0.497 | 0.443 (0.372) | 1.19 | 1.447 (0.557) | **2.596 *** |
| Floral stems^2 | -0.497 (0.297) | -1.675 | 0.418 (0.461) | 0.905 | -0.482 (0.384) | -1.256 | -0.862 (0.505) | -1.708 |
| Flowers^2 | 0.342 (0.231) | 1.482 | -0.166 (0.321) | -0.516 | 0.438 (0.298) | 1.47 | 0.368 (0.336) | 1.094 |
| Dry mass^2 | -0.649 (0.192) | **-3.376 ***** | -0.125 (0.315) | -0.397 | -0.265 (0.248) | -1.068 | 0.771 (0.364) | **2.12 *** |

| **(b) *M.* x *robertsii*** | **Stolons** | | | |
| --- | --- | --- | --- | --- |
| | **IOW** | | **SHE** | |
| | **Estimate (SE)** | ***t*** | **Estimate (SE)** | ***t*** |
| Intercept | 0.787 (0.058) | 13.506 *** | 0.842 (0.058) | 14.393 *** |
| Height | 0.164 (0.314) | 0.522 | -0.258 (0.298) | -0.865 |
| Stomata | -0.264 (0.362) | -0.73 | -0.506 (0.362) | -1.396 |
| Flowering day | -0.229 (0.383) | -0.598 | 0.58 (0.754) | 0.77 |
| Flowering node | -0.342 (0.389) | -0.879 | -0.478 (0.348) | -1.373 |
| Corolla | 0.725 (0.548) | 1.324 | 0.828 (0.317) | **2.608 *** |
| Plant cover | 0.164 (0.381) | 0.429 | 0.196 (0.493) | 0.397 |
| Branches | 0.236 (0.219) | 1.079 | 0.204 (0.426) | 0.478 |
| Floral stems | 0.24 (0.265) | 0.907 | -0.196 (0.403) | -0.488 |
| Flowers | -0.287 (0.303) | -0.949 | 0.039 (0.33) | 0.117 |
| Dry mass | 0.615 (0.297) | **2.073 *** | -0.385 (0.424) | -0.907 |
| Height^2 | -0.241 (0.595) | -0.405 | 0.222 (0.61) | 0.364 |
| Stomata^2 | 0.532 (0.71) | 0.75 | 1.075 (0.722) | 1.488 |
| Flowering day^2 | 0.74 (0.788) | 0.939 | -0.85 (1.498) | -0.567 |
| Flowering node^2 | 0.293 (0.779) | 0.376 | 0.842 (0.678) | 1.242 |
| Corolla^2 | -1.354 (1.09) | -1.242 | -1.519 (0.635) | **-2.39 *** |
| Plant cover^2 | -0.594 (0.635) | -0.936 | -0.121 (0.853) | -0.142 |
| Branches^2 | -0.286 (0.352) | -0.812 | -0.171 (1.005) | -0.17 |
| Floral stems^2 | -0.801 (0.483) | -1.659 | 0.081 (0.91) | 0.089 |
| Flowers^2 | 1.071 (0.571) | 1.877 | -0.005 (0.602) | -0.009 |
| Dry mass^2 | -0.758 (0.504) | -1.504 | 0.433 (0.705) | 0.614 |

| (c) *M. luteus* | Fruits | | | | Stolons | | | |
|---|---|---|---|---|---|---|---|---|
| | IOW | | SHE | | IOW | | SHE | |
| | **Estimate (SE)** | ***t*** | **Estimate (SE)** | ***t*** | **Estimate (SE)** | ***t*** | **Estimate (SE)** | ***t*** |
| Intercept | 0.941 (0.047) | 19.872 *** | 0.864 (0.074) | 11.696 *** | 0.838 (0.055) | 15.201 *** | 0.831 (0.115) | 7.213 *** |
| Height | -0.158 (0.486) | -0.325 | 0.354 (0.542) | 0.653 | 0.987 (0.565) | 1.747 | -0.19 (0.892) | -0.213 |
| Stomata | -0.788 (0.39) | -2.023 | -0.154 (0.562) | -0.274 | 1.012 (0.453) | **2.232 *** | -0.707 (0.917) | -0.771 |
| Flowering day | -0.11 (0.388) | -0.284 | -0.611 (0.976) | -0.625 | 0.576 (0.451) | 1.275 | 1.936 (1.361) | 1.423 |
| Flowering node | -0.381 (0.333) | -1.147 | -0.098 (0.417) | -0.234 | 0.336 (0.387) | 0.868 | -0.44 (0.679) | -0.647 |
| Corolla | 1.34 (0.666) | 2.013 | 1.18 (1.002) | 1.178 | -1.249 (0.775) | -1.613 | -0.209 (1.572) | -0.133 |
| Plant cover | -0.529 (0.347) | -1.526 | -0.183 (0.426) | -0.429 | 0.08 (0.404) | 0.197 | 0.566 (0.694) | 0.816 |
| Branches | 0.524 (0.319) | 1.641 | 0.093 (0.479) | 0.194 | -0.422 (0.372) | -1.137 | 0.274 (0.684) | 0.401 |
| Floral stems | -0.337 (0.361) | -0.933 | -0.466 (0.48) | -0.971 | -0.286 (0.42) | -0.682 | -0.087 (0.76) | -0.114 |
| Flowers | 0.04 (0.318) | 0.127 | 0.88 (0.335) | **2.632 *** | 0.088 (0.37) | 0.239 | 0.594 (0.549) | 1.082 |
| Dry mass | 0.86 (0.307) | **2.798 **** | -0.098 (0.387) | -0.253 | -0.492 (0.358) | -1.375 | -0.668 (0.625) | -1.067 |
| Height^2 | 0.203 (0.936) | 0.216 | -0.372 (1.036) | -0.359 | -1.681 (1.09) | -1.543 | 0.451 (1.712) | 0.264 |
| Stomata^2 | 1.592 (0.729) | **2.185 *** | 0.281 (1.114) | 0.253 | -2.07 (0.848) | **-2.442 *** | 1.234 (1.819) | 0.679 |
| Flowering day^2 | 0.042 (0.618) | 0.067 | 1.585 (1.999) | 0.793 | -0.703 (0.72) | -0.978 | -3.918 (2.743) | -1.428 |
| Flowering node^2 | 0.371 (0.648) | 0.573 | 0.133 (0.805) | 0.166 | -0.923 (0.754) | -1.224 | 1.261 (1.288) | 0.979 |
| Corolla^2 | -2.294 (1.278) | -1.794 | -2.222 (1.993) | -1.115 | 2.142 (1.488) | 1.44 | 0.508 (3.116) | 0.163 |
| Plant cover^2 | 1.001 (0.648) | 1.545 | 0.576 (0.758) | 0.76 | 0.092 (0.754) | 0.122 | -1.443 (1.205) | -1.197 |
| Branches^2 | -1.483 (0.684) | **-2.169 *** | 0.18 (0.944) | 0.191 | 0.744 (0.795) | 0.935 | -0.288 (1.191) | -0.242 |
| Floral stems^2 | 1.199 (0.696) | 1.723 | 0.215 (0.831) | 0.259 | 0.524 (0.81) | 0.647 | -0.338 (1.266) | -0.267 |
| Flowers^2 | 0.042 (0.652) | 0.065 | -1.077 (0.549) | -1.962 | -0.264 (0.758) | -0.348 | -0.638 (0.913) | -0.699 |
| Dry mass^2 | -1.484 (0.573) | **-2.59 *** | 0.048 (0.773) | 0.062 | 0.692 (0.667) | 1.038 | 1.23 (1.261) | 0.975 |

**Table note:** * P<0.05; ** P<0.01; *** P<0.001.

**Table S9**. Results of the GLMMs modelling the effects of experimental site (S), species (Sps) and their interaction in 12 fitness, phenotypic and phenological traits recorded in the only existing introduced population of *M .luteus*, and in *M. guttatus* and *M.* × *robertsii* populations from the north range of UK, during the reciprocal transplants experiment. The models comprising *M. luteus* and north populations of the other species include *M. guttatus* for fruits and stolons and *M.* x *robertsii* for stolons and the rest of phenotypic traits.

| | | ***M. luteus* + north populations** | | ***M. luteus*** | |
|---|---|---|---|---|---|
| **Trait** | **Fixed factor** | **Estimate (SE)** | **$\chi^2$** | **Estimate (SE)** | **$\chi^2$** |
| **Fruits** | Intercept | 2.855 (0.310) | 84.708 *** | 3.028 (0.244) | 153.422 *** |
| | **$W_0$** | -0.001 (0.000) | 1.931 | 0.001 (0.002) | 0.37 |
| | **Site** (SHE) | -0.632 (0.040) | **248.94 *** ** | -0.432 (0.044) | **96.963 *** ** |
| | **Sps** (*luteus*) | 0.310 (0.996) | 0.097 | | |
| | **Sps:Site** (*luteus*:SHE) | 0.188 (0.056) | **11.018 *** ** | | |
| **Stolons** (guttatus) | Intercept | 1.978 (0.161) | 151.243 | 2.169 (0.173) | 156.919 *** |
| | **$W_0$** | -0.002 (0.001) | 2.17 | -0.006 (0.004) | 2.08 |
| | **Site** (SHE) | -0.01 (0.06) | 0.023 | 0.132 (0.072) | 3.33 |
| | **Sps** (*robertsii*) | 0.041 (0.165) | 0.061 | | |
| | **Sps:Site** (*robertsii*:SHE) | 0.156 (0.093) | 2.797 | | |
| **Stolons** (robertsii) | Intercept | 1.949 (0.109) | 316.918 *** | | |
| | **$W_0$** | 0 (0) | 0.291 | | |
| | **Site** (SHE) | 0.155 (0.071) | 4.825 * | | |
| | **Sps** (*robertsii*) | -0.5 (0.149) | **11.333 *** ** | | |
| | **Sps:Site** (*robertsii*:SHE) | 0.304 (0.101) | **9.078 ** ** | | |
| **Flowering day** | Intercept | 41.764 (5.172) | 65.216 *** | 50.292 (4.257) | 139.552 *** |
| | **$W_0$** | -0.009 (0.007) | 1.691 | -0.231 (0.103) | 4.986 * |
| | **Site** (SHE) | 20.916 (1.829) | **130.82 *** ** | 19.92 (1.821) | **119.599 *** ** |
| | **Sps** (*robertsii*) | -3.656 (5.632) | 0.421 | | |
| | **Sps:Site** (*robertsii*:SHE) | 4.475 (2.362) | 3.589 | | |
| **Flowering node** | Intercept | 5.882 (0.52) | 128.041 *** | 6.896 (0.789) | 76.434 *** |
| | **$W_0$** | 0 (0.001) | 0.081 | -0.026 (0.019) | 1.858 |
| | **Site** (SHE) | -0.337 (0.267) | 1.593 | -0.459 (0.344) | 1.783 |
| | **Sps** (*robertsii*) | -1.413 (0.585) | 5.835 * | | |
| | **Sps:Site** (*robertsii*:SHE) | 0.465 (0.346) | 1.807 | | |

| | | | | | |
|---|---|---|---|---|---|
| **Branches** | Intercept | 1.989 (0.095) | 442.485 *** | 1.735 (0.177) | 96.115 *** |
| | **$W_0$** | 0.001 (0) | 5.914 * | 0.007 (0.004) | 3.596 |
| | **Site** (SHE) | 0.202 (0.067) | **8.973 **** | 0.233 (0.07) | **11.069 ***** |
| | **Sps** (*robertsii*) | 0.157 (0.123) | 1.642 | | |
| | **Sps:Site** (*robertsii*:SHE) | -0.272 (0.088) | **9.596 **** | | |
| **Stems** | Intercept | 2.427 (0.103) | 550.925 *** | 2.185 (0.165) | 175.731 *** |
| | **$W_0$** | 0.001 (0) | **17.841 ***** | 0.007 (0.003) | 5.628 * |
| | **Site** (SHE) | 0.131 (0.053) | 5.976 * | 0.159 (0.055) | **8.251 **** |
| | **Sps** (*robertsii*) | 0.184 (0.134) | 1.878 | | |
| | **Sps:Site** (*robertsii*:SHE) | -0.254 (0.07) | **13.314 ***** | | |
| **Flowers** | Intercept | 4.948 (0.083) | 3511.655 *** | 4.81 (0.118) | 1654.504 *** |
| | **$W_0$** | 0 (0) | **56.073 ***** | 0.004 (0.001) | **20.025 ***** |
| | **Site** (SHE) | -0.043 (0.016) | 6.993 ** | -0.029 (0.016) | 3.007 |
| | **Sps** (*robertsii*) | 0.161 (0.109) | 2.194 | | |
| | **Sps:Site** (*robertsii*:SHE) | -0.101 (0.021) | **22.965 ***** | | |
| **Corolla** | Intercept | 332.43 (32.097) | 107.27 *** | 790.771 (95.861) | 413.439 *** |
| | **$W_0$** | -0.071 (0.041) | 3.001 | -1.474 (2.309) | 6.956 ** |
| | **Site** (SHE) | 7.133 (10.831) | 0.434 | -69.603 (44.065) | 0.076 |
| | **Sps** (*robertsii*) | 89.191 (34.911) | 6.527 * | | |
| | **Sps:Site** (*robertsii*:SHE) | 6.448 (14.033) | 0.211 | | |
| **Stomata** | Intercept | 16.28 (2.911) | 31.252 *** | 18.028 (1.263) | 203.76 *** |
| | **$W_0$** | 0.001 (0.002) | 0.048 | -0.046 (0.03) | 2.362 |
| | **Site** (SHE) | -2.036 (0.6) | **11.538 ***** | -2.254 (0.561) | **16.127 ***** |
| | **Sps** (*robertsii*) | -1.842 (3.127) | 0.347 | | |
| | **Sps:Site** (*robertsii*:SHE) | 1.835 (0.791) | 5.378 * | | |
| **Height** | Intercept | 22.48 (1.478) | 231.404 *** | 19.889 (3.452) | 33.197 *** |
| | **$W_0$** | 0.024 (0.006) | **14.156 ***** | 0.092 (0.082) | 1.277 |
| | **Site** (SHE) | 1.316 (1.539) | 0.731 | 1.623 (1.541) | 1.109 |
| | **Sps** (*robertsii*) | -0.809 (1.957) | 0.171 | | |
| | **Sps:Site** (*robertsii*:SHE) | -1.616 (2.028) | 0.635 | | |
| **Cover** | Intercept | 716.387 (113.072) | 40.141 *** | 790.771 (95.861) | 68.048 *** |
| | **$W_0$** | 0.496 (0.171) | **8.422 **** | -1.474 (2.309) | 0.407 |
| | **Site** (SHE) | -60.482 (41.944) | 2.079 | -69.603 (44.065) | 2.495 |
| | **Sps** (*robertsii*) | -85.94 (124.445) | 0.477 | | |
| | **Sps:Site** (*robertsii*:SHE) | 1.896 (55.233) | 0.001 | | |
| **Dry mass** | Intercept | 37.018 (6.576) | 31.692 *** | 21.856 (5.162) | 17.93 *** |
| | **$W_0$** | 0.021 (0.009) | 4.989 * | 0.421 (0.126) | **11.132 ***** |
| | **Site** (SHE) | -8.97 (2.304) | **15.157 ***** | -7.106 (2.476) | **8.234 **** |
| | **Sps** (*robertsii*) | -1.899 (7.197) | 0.07 | | |
| | **Sps:Site** (*robertsii*:SHE) | -0.73 (3.039) | 0.058 | | |

**Table note:** Maximal model Estimates and Standard Errors for each fixed factor and interaction are provided jointly with results of the type-III Wald $\chi$2 tests. $\chi$2 values and indications of their associated *P*-values are provided. * $P < 0.05$; ** $P < 0.01$; *** $P < 0.001$. Significant effects after Bonferroni correction of *P*-values are indicated in bold. $\chi$2 degrees of freedom=1-2.

**Figure S1.** Temperature and photoperiod conditions of the environmental models included in the chambers experiment. Squares represent day temperature and circles represent number of light hours per day in each fortnight in each of four chambers. Orange and blue symbols represent natural conditions in Isle of Wight and Shetland, respectively. Treatment codes indicate photoperiod (L, long; S, short) and temperature (W, warm; C, cold).

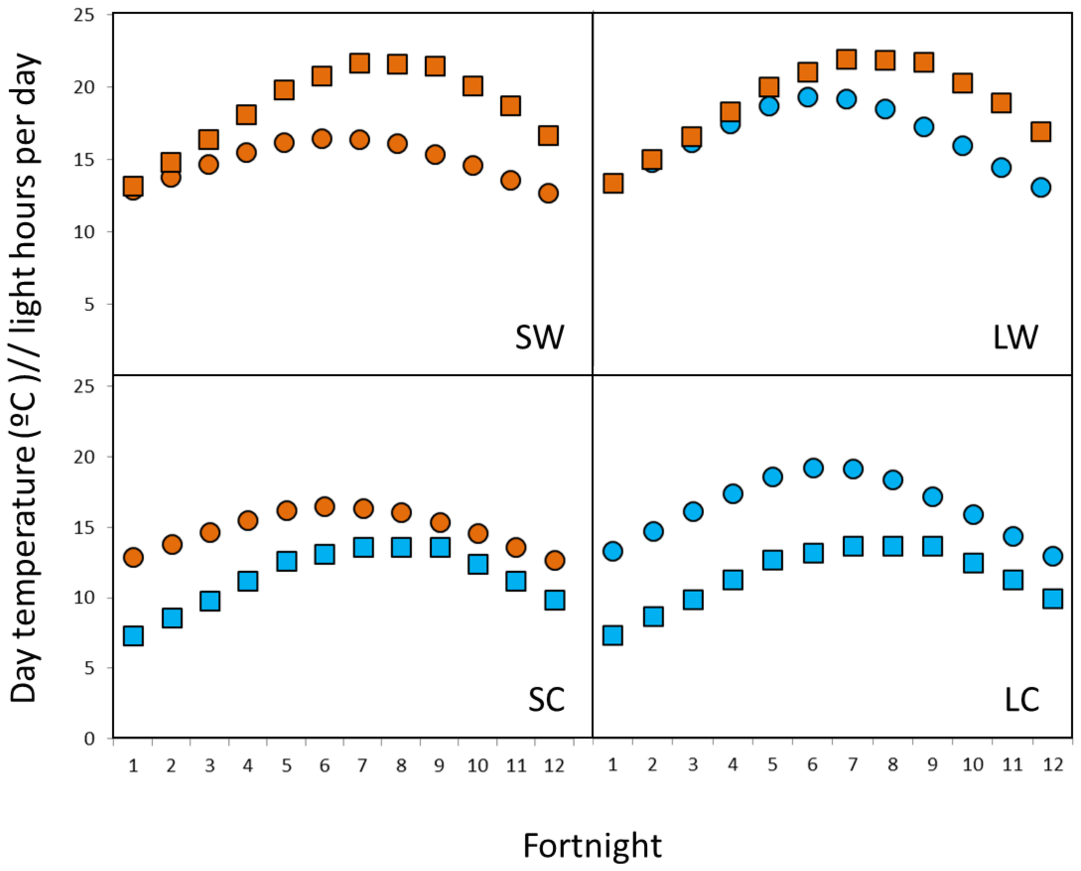

**Figure S2.** Correlation (A) and Principal Component Analyses (B) of the phenotypic traits measured in the controlled environmental chambers experiments for native *M. guttatus*, invasive *M. guttatus*, and *M*. x *robertsii* populations, and the three population types together.

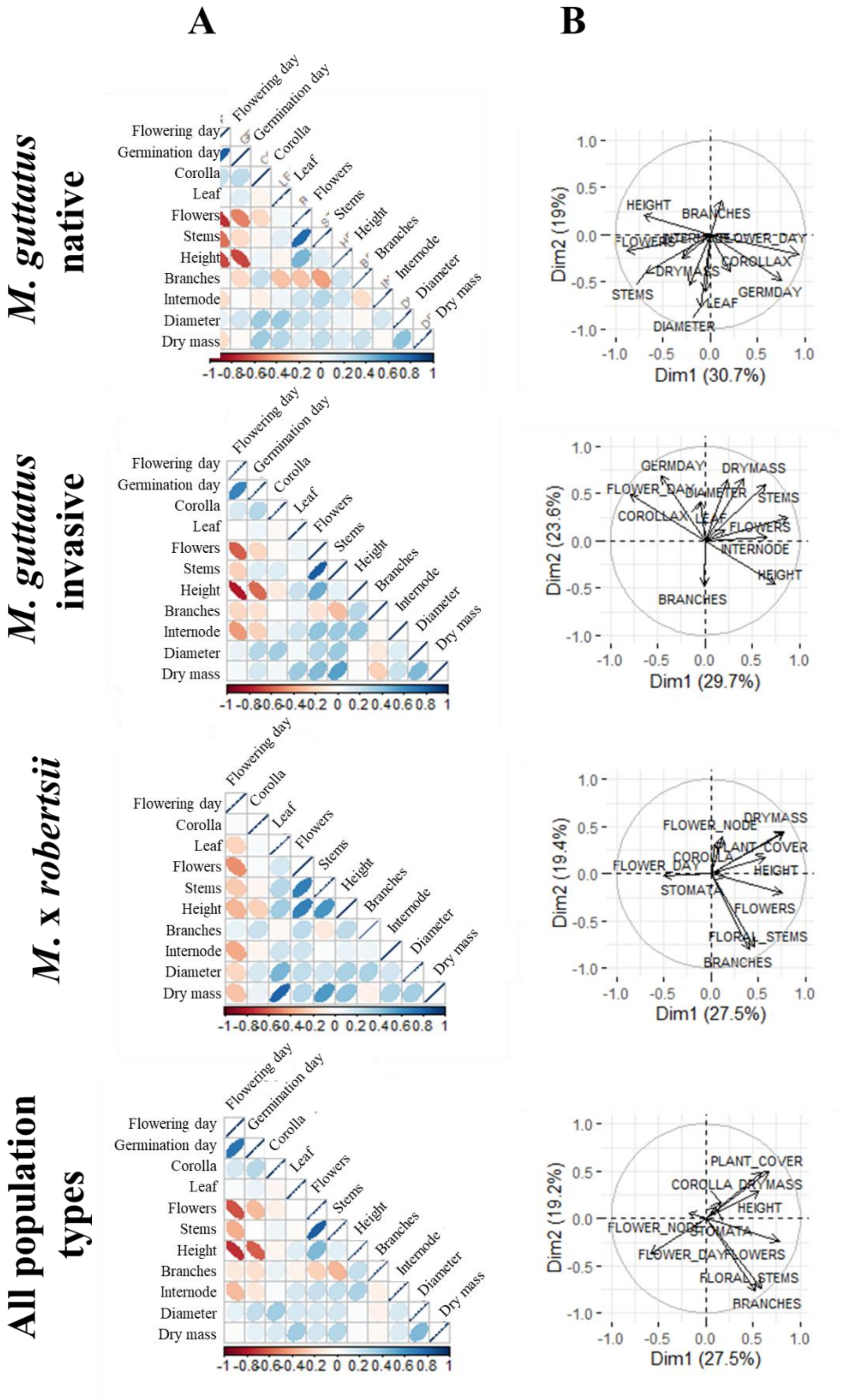

**Figure S3.** Correlation (A) and Principal Component Analyses (B) of the phenotypic traits measured for phenotypic selection analyses in the reciprocal transplants experiments for *M. guttatus*, *M.* x *robertsii*, *M. luteus*, and the three species togheter.

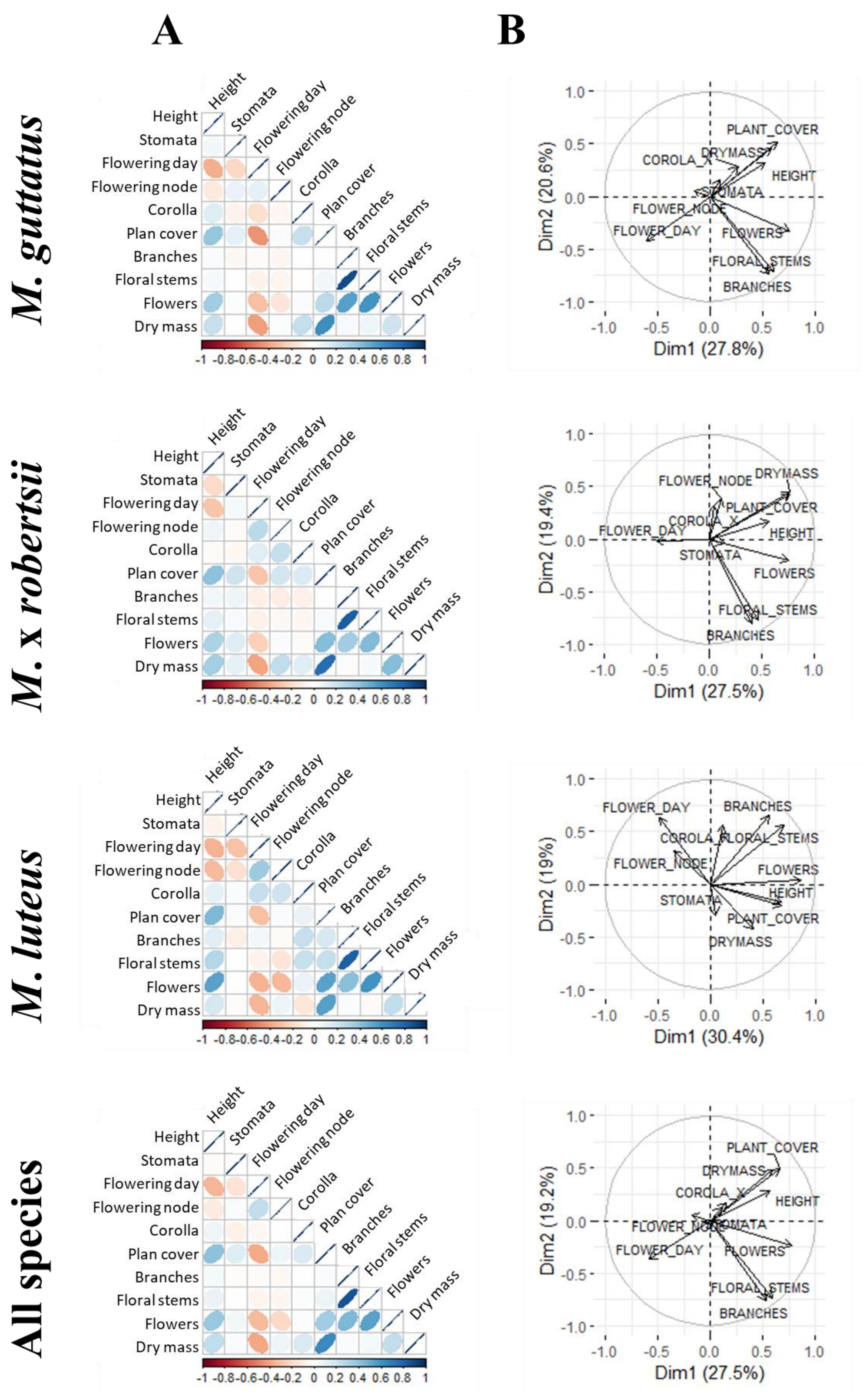

**Figure S4.** Phenotypic traits of native *Mimulus guttatus* (US), introduced *M. guttatus* (UK), and *M.* × *robertsii* (ROB) grew in four different controlled environmental chambers with contrasting photoperiods (L: long; S: short) and temperatures (W: warm; C: cold) in a crossed design. Mean values and standard errors of the variables measured are indicated by dots and error bars, respectively. Units as follows: Corolla width (mm), node diameter (mm), plant height (cm), internode length (cm), dry mass (g).

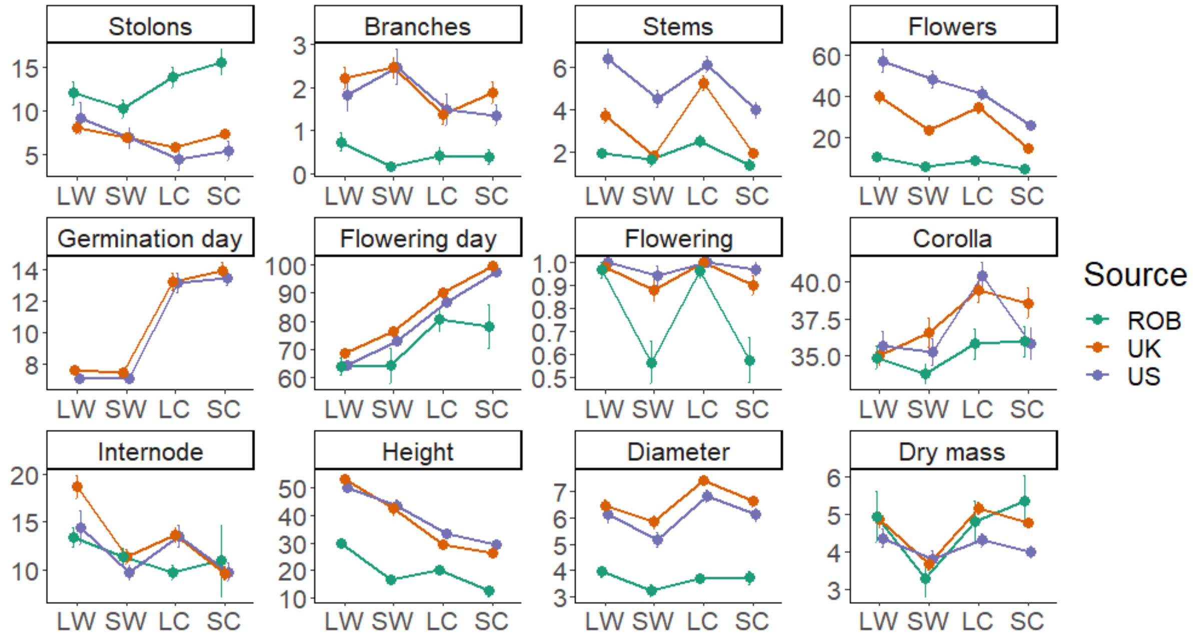

**Figure S5.** Reaction norms for each variable measured in the reciprocal transplant experiment of populations of *Mimulus guttatus* and *M.* × *robertsii* from different latitudes in the British Isles. Mean values and standard errors of the variables measured are indicated by dots and error bars, respectively. Units as follows: Corolla width (mm), plant height (cm), internode length (cm), dry mass (g), stomata (number per cm$^2$), plant cover (cm$^2$).

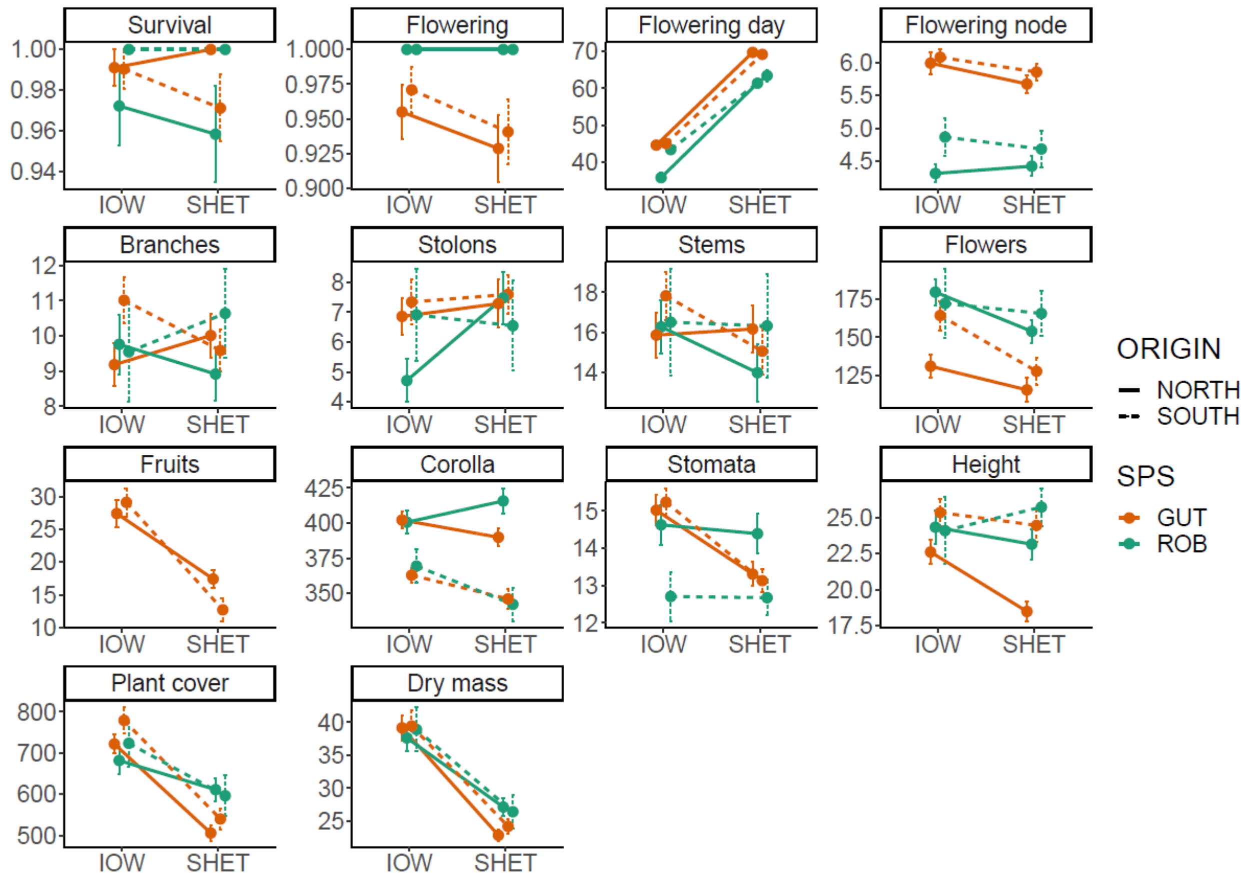

**Figure S6.** (A) Sexual and asexual fitness (mean number of fruits or stolons ± s.d.) of the *M. luteus* individuals from the single population of this species included in the transplants experiment at different latitudes in the UK. (B) Estimates and 95% confidence intervals for the phenotypic selection coefficients on each trait and site included in the selection gradients.

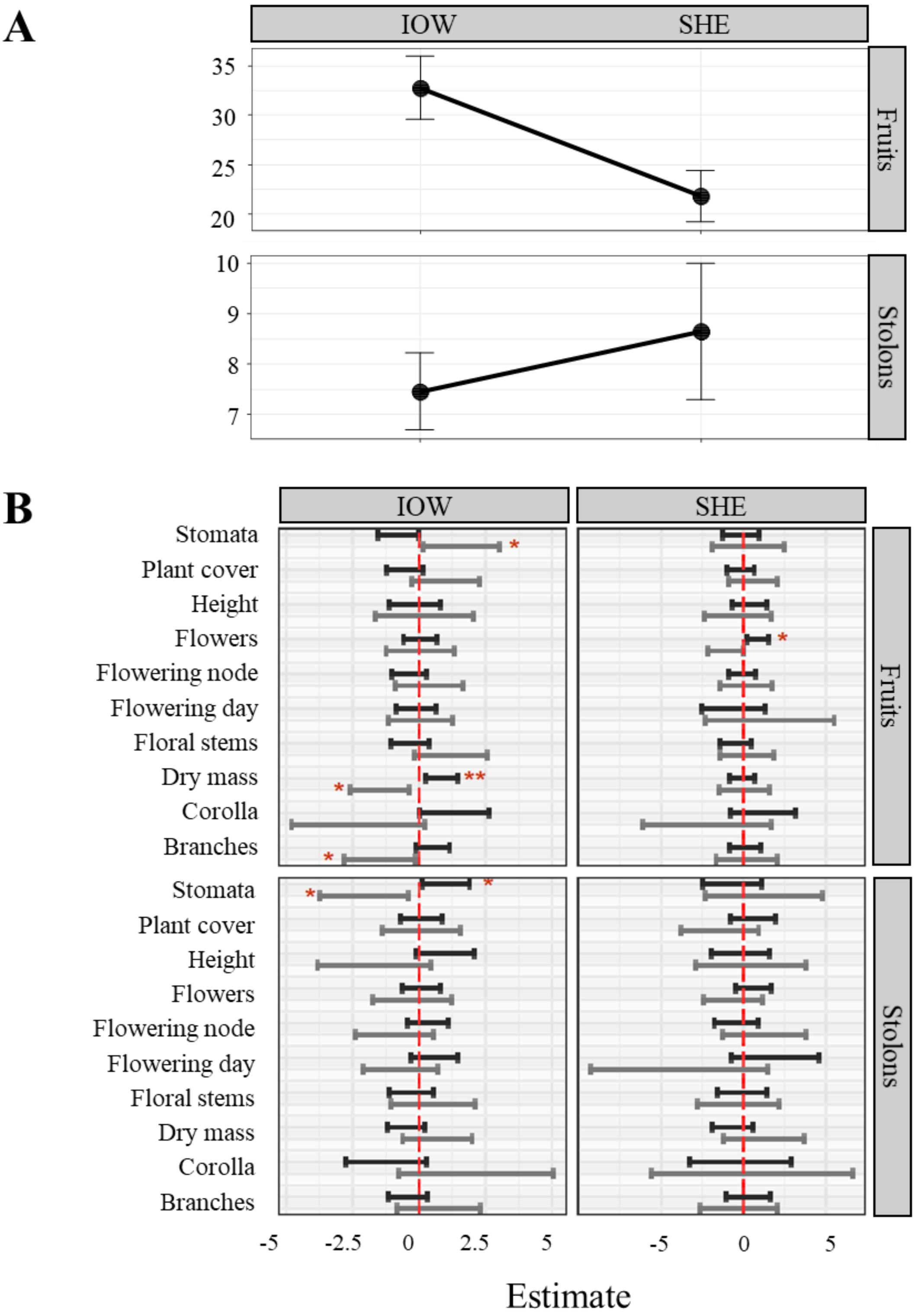
